# Bosutinib stimulates macrophage survival, phagocytosis and intracellular killing of bacteria

**DOI:** 10.1101/2023.12.13.571434

**Authors:** Ronni A. G. da Silva, Claudia J. Stocks, Guangan Hu, Kimberly A. Kline, Jianzhu Chen

## Abstract

Host-acting compounds are emerging as potential alternatives to combat antibiotic resistance. Here, we show that bosutinib, an FDA-approved chemotherapeutic for treating chronic myelogenous leukemia, does not possess any antibiotic activity but enhances macrophage responses to bacterial infection. *In vitro*, bosutinib stimulates murine and human macrophages to kill bacteria more effectively. In a murine wound infection with vancomycin-resistant *Enterococcus faecalis*, a single intraperitoneal bosutinib injection or multiple topical applications on the wound reduces bacterial load by approximately 10-fold, which is abolished by macrophage depletion. Mechanistically, bosutinib stimulates macrophage phagocytosis of bacteria by upregulating surface expression of bacterial uptake markers Dectin-1 and CD14 and promoting actin remodelling. Bosutinib also stimulates bacterial killing by elevating the intracellular levels of reactive oxygen species. Moreover, bosutinib drives NF-κB activation which protects infected macrophages from dying. Other Src kinase inhibitors such as DMAT and Tirbanibulin also upregulate expression of bacterial uptake markers in macrophages and enhance intracellular bacterial killing. Finally, co-treatment with bosutinib and mitoxantrone, another chemotherapeutic in clinical use, results in an additive effect on bacterial clearance *in vitro* and *in vivo*. These results show that bosutinib stimulates macrophage clearance of bacterial infection through multiple mechanisms and could be used to boost host innate immunity to combat drug-resistant bacterial infections.

**Significance:** This study shows that bosutinib, an FDA-approved chemotherapeutic, stimulates macrophage responses to antibiotic-resistant bacterial infection by enhancing phagocytosis and intracellular killing of bacteria and promoting survival of infected macrophages. These findings suggest that bosutinib could serve as an adjuvant therapy to combat drug resistant bacterial infections and opens the possibility to target Src kinases to boost innate immunity in general.

## Introduction

Antimicrobial resistance greatly limits the treatment options for bacterial infections. Compounds that enhance the host immune responses are emerging as alternative approaches to treat antibiotic resistant infections [1]. However, since many bacteria have evolved mechanisms to escape or suppress the host immune responses either extracellularly or intracellularly [2, 3], any adjuvant therapy should ideally enhance both bacterial uptake and intracellular killing for optimal clearance of infection.

Macrophages play a critical role in defence against bacterial infection by recognizing and phagocytosing bacteria and killing them intracellularly in phagolysosomes. Bacterial recognition is mediated by an array of receptors that recognize evolutionarily conserved pathogen-associated molecular patterns (PAMPs). Binding of receptors to PAMPs triggers a signalling cascade that facilitates the process of phagocytosis [4]. Specifically, engagement of phagocytic receptors triggers signalling pathways that prompt a reorganization of the actin cytoskeleton and membrane lipids [5], leading to membrane expansion and engulfment of bacteria. Once internalised, sequential intracellular trafficking events that involve fusion and fission with endocytic vesicles and the lysosome result in the formation of the anti-microbial phagolysosome [6, 7]. The phagolysosome deploys different mechanisms to kill and degrade bacteria, including low pH, reactive oxygen species (ROS), and hydrolytic enzymes.

The Src kinase family (SFK) consists of nine non-receptor tyrosine kinases in mammals: SRC, LCK, LYN, BLK, HCK, FYN, FGR, YES, and YRK [8]. These kinases function in many cellular processes including cell adhesion and migration, proliferation, differentiation, apoptosis and metabolism [9–12]. SFK members play a crucial role in host defence and inflammation, mediating signalling from cell surface receptors in hematopoietic cells and orchestrating adhesion and transmigration during leukocyte recruitment [13, 14]. Modulation of SRC kinase activity has been investigated for its chemotherapeutic potential [15] and bosutinib was developed to treat chronic myelogenous leukemia by inhibiting SRC and ABL kinases. SRC kinase activation has been shown to contribute to innate immune responses to viral infections [16] and SRC kinase inhibition is known to prevent the assembly of dengue virions and ameliorate sepsis outcomes in a murine model of polymicrobial sepsis [17, 18]. In macrophages, SRC kinase activity contributes to adhesion, migration, and phagocytosis [19], suggesting that SRC kinase activity is important for innate immune responses to infections.

We have previously identified small molecule compounds that enhance the ability of macrophages to clear bacterial infection, including bosutinib (BOS), which stimulates macrophage intracellular killing of *E. faecalis*, *Salmonella Typhimurium,* and uropathogenic *Escherichia coli in vitro* [1]. BOS is an orally available SRC-ABL tyrosine kinase inhibitor used to treat chronic myelogenous leukaemia [20] by inhibiting tumor cell proliferation [21]. In this study, we show BOS treatment reduces the bacterial burden in a murine wound model of infection with a vancomycin-resistant strain of *E. faecalis* (VRE) or methicillin-resistant strain of *S. aureus* (MRSA) in a macrophage-dependent manner. We have also elucidated the mechanisms by which BOS enhance macrophage responses to bacterial infection. We show that BOS inhibition of SRC kinase activity affects SLK phosphorylation, leading to upregulation of bacterial uptake surface markers and actin-remodelling and therefore enhanced phagocytic activity of macrophages. BOS also stimulates macrophages to kill phagocytosed bacteria by upregulating ROS production and survival of infected macrophages. These findings suggest the potential for repurposing of BOS as an immune boosting adjunct therapy for treating antibiotic resistant bacterial infections.

## Results

### BOS enhances macrophage clearance of bacteria *in vitro* and *in vivo*

We previously showed that BOS stimulates intracellular killing of the *E. faecalis* (strain OG1RF) by murine macrophage cell line RAW264.7 [22]. Here, we further tested the ability of BOS to stimulate killing of vancomycin-resistant *E. faecalis* strain V583 (VRE) by RAW264.7, murine bone marrow-derived macrophages (BMDM), the human monocytic cell line THP-1, and human monocyte-derived macrophages (HMDM). In these assays, cells were infected with VRE for 3h and non-attached VRE were removed by washing. Residual extracellular bacteria were eliminated by the addition of gentamicin and penicillin in the culture medium for the entire duration of the assay. Simultaneous to antibiotic addition, the cultures were treated with BOS (0.52 μg/mL) or without BOS and the number of intracellular VRE was quantified 15 h later. As shown in **Fig. 1A**, BOS treatment resulted in a statistically significant reduction of intracellular CFU by ∼1-log in RAW264.7 cells and ∼0.5-log in THP-1 cells, but a non-significant reduction in BMDM and HMDM. Similarly, BOS treatment also stimulated macrophage killing of other intracellular bacterial species, including MRSA, *P. aeruginosa* and the multi-drug resistant *E. coli* strain 958 (**Fig. S1A**). When VRE, MRSA, *P. aeruginosa* and *E. coli* EC958 were incubated with increasing concentrations of BOS in the absence of host cells, the minimum inhibitory concentration (MIC) was greater than 13 μg/mL, which was 25-fold higher than the 0.52 μg/mL BOS used to treat macrophages **(Table S1 and S2)**, suggesting that BOS does not have direct antibiotic activity. Consistently, when RAW264.7 cells or BMDM were pretreated with BOS for 18 h prior to VRE infection, the levels of intracellular CFU were reduced by ∼1-log and ∼0.5-log, respectively, compared to the untreated controls (**Fig. 1B**). Together these results show that BOS does not possess antibiotic activity but can stimulate macrophages to more effectively eliminate intracellular bacteria.

**Figure 1.**
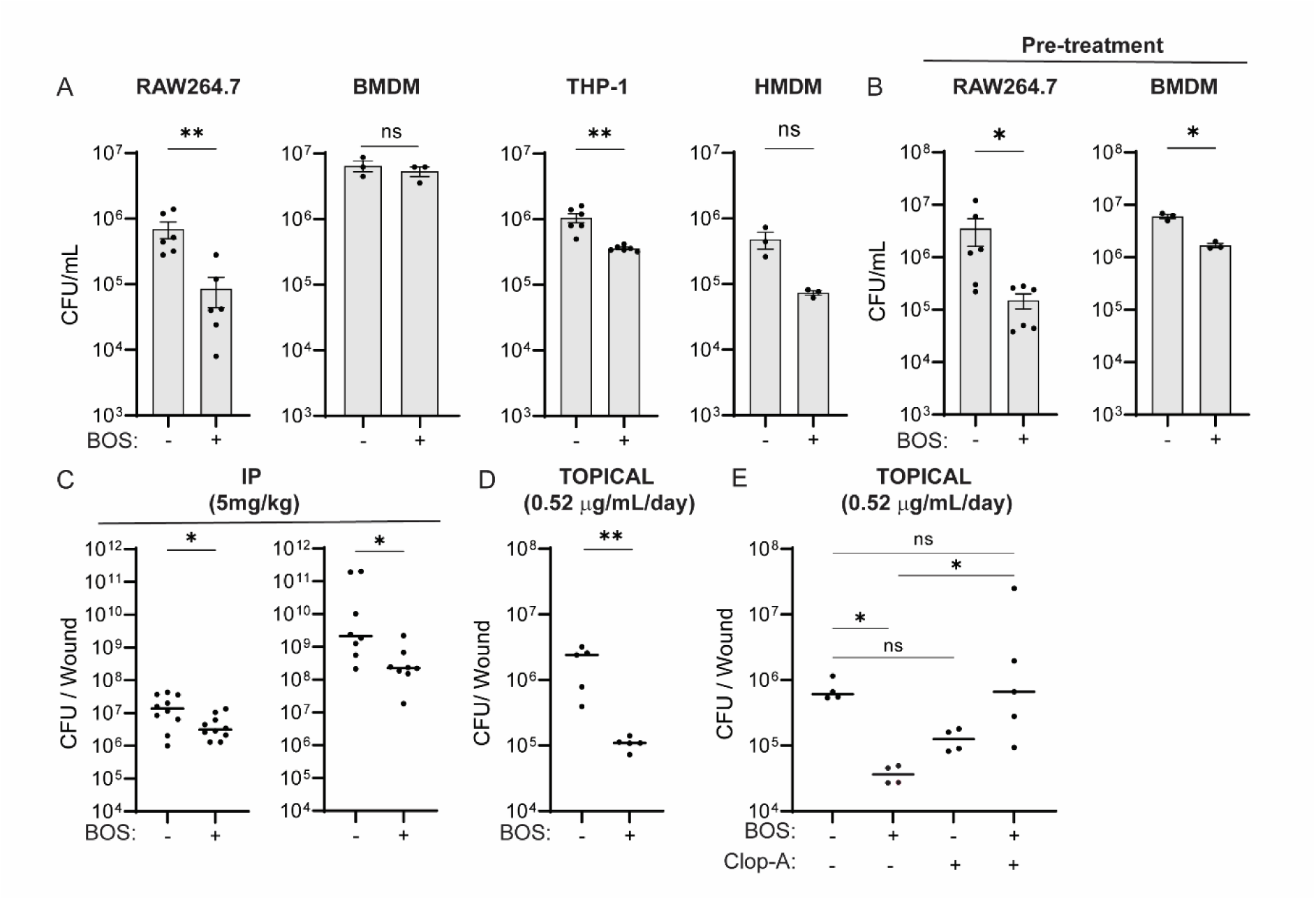
- BOS enhances macrophage killing of intracellular bacteria *in vitro* and *in vivo*. **(A)** Comparison of VRE CFU in RAW264.7, BMDM, THP-1, and HMDM cells treated with BOS for 15 h after initial infection of 3 h (0.52 μg/mL). **(B)** Comparison of VRE CFU in overnight BOS-pretreated RAW264.7 and BMDM cells. Data shown (mean ± SEM) is summary of at least three independent experiments (A-B). **(C)** Comparison of VRE (left) or MRSA (right) CFU per infected wound from animals treated with a single IP injection of either vehicle (DMSO) or BOS (5 mg/kg in 30 μL DMSO). **(D)** Comparison of VRE CFU per infected wound treated with five topical doses of vehicle (PBS) or BOS (0. 52 μg/ml). **(E)** Comparison of VRE CFU per wound treated with five topical doses of vehicle or BOS with or without macrophage depletion with Clop-A. Each symbol represents one mouse with the median indicated by the horizontal line. Data were from two independent experiments with two to five mice per experiment. Statistical analysis was performed using unpaired t test with Welch’s corrections (A-B), the nonparametric Mann-Whitney test to compare ranks (C-D), and ordinary one-way ANOVA, followed by Tukey’s multiple comparison test (E). For all analyses, NS denotes not significant; *P ≤ 0.05 and **P ≤ 0.01.

We evaluated the effect of BOS *in vivo* using a murine wound infection model with VRE and MRSA. Wounds were infected with 10^6^ colony-forming units (CFU) of VRE or MRSA, and were simultaneously given a single dose intraperitoneally (IP) of BOS (5 mg/kg in 30 μL) or vehicle (DMSO) at the time of infection. Twenty-four hours post infection (hpi), wounds were excised and CFU of VRE and MRSA were enumerated. BOS treatment resulted in a reduction of VRE and MRSA CFU by 0.6-log and 1.2-log, respectively, compared to vehicle-treated wounds (**Fig. 1C**). When five doses of BOS were given IP, VRE CFU were reduced by 1.2-log **(Fig. S1B)**. Alternatively, when infected wounds were treated topically with a daily dose of 10μL PBS containing 0.52 μg/ml of BOS for 5 days, VRE CFU were reduced by ∼1.3-log as compared to PBS treatment (**Fig. 1D**). Topical treatments of infected wounds with BOS resulted in wound diameters that were half the size of vehicle-treated wounds, which correlated with reduced bacterial burden **(Fig. S1C)**. Furthermore, mice pretreated with a single IP dose of BOS (5 mg/kg) 24h prior to infection had ∼0.7-log fewer VRE CFU in wounds at the end of the experiment as compared to vehicle-treated animals **(Fig. S1D)**, whereas a single topical dose of BOS, even at 10x higher concentration (5.2 μg/ml), did not result in a significant reduction of the bacterial burden in the wounds **(Fig. S2E)**. Thus, BOS stimulates bacterial clearance *in vivo*.

To verify the requirement for macrophages in BOS-stimulated bacterial clearance *in vivo*, we depleted macrophages by using liposomes containing clodronate (clophosome-A (clop-A)) [23–25]. Mice were injected IP with clop-A (200 μL, 6 mg/mL) 3 days prior to wounding and infection, with additional doses of clop-A on the day of wounding and infection and every 2 days afterwards. In addition, clop-A (10 μL, 6 mg/mL) was applied to the wounds every 2 days **(Fig. S1F)**. Empty liposomes were used as vehicle control. Following VRE infection, BOS (10 μL, 0.52 μg/ml) was applied to the wounds daily for 5 days as above (**Fig. 1D**). Five days after wounding and infection, mice were sacrificed, and wounds were excised and dissociated for macrophage and CFU analysis. Among CD45^+^ leukocytes, the percentages of CD11b^+^ F480^+^ macrophages, but not Ly6G^+^ neutrophils, were reduced from ∼55% in vehicle treated mice to ∼1.5 % in clop-A treated mice, regardless of BOS treatment, suggesting successful depletion of macrophages from the wounds **(Fig. S1G-I**). Without depletion of macrophages, BOS treatment reduced VRE CFU by ∼1.3-log (**Fig. 1E**). By contrast, with macrophage depletion, BOS did not significantly reduce VRE CFU in the wounds as compared to either no macrophage depletion or PBS-treated macrophage-depleted wounds (**Fig. 1E**). These results show that BOS- stimulated clearance of bacterial infection *in vivo* is primarily mediated by macrophages.

### BOS stimulates macrophage phagocytosis of bacteria through actin-remodelling

To elucidate the mechanisms by which BOS stimulates macrophage clearance of bacteria, we first tested if BOS stimulates macrophage phagocytosis of bacteria. RAW264.7 were pre-treated with BOS for 18 h and then incubated with Syto9-stained VRE for 30 min. Uptake of fluorescent bacteria by macrophages was measured after quenching extracellular bacterial fluorescence by trypan blue. Fluorescence intensity was two times higher in RAW264.7 cells that were pre-treated with BOS than non-treated cells (**Fig. 2A**), indicating phagocytosis of more VRE. Intracellular CFU were also directly measured following 3 h of infection, washing, and antibiotic inhibition of the extracellular bacteria for 30 min as described above. VRE CFU were 0.5-log higher in BOS-pretreated macrophages than non-treated cells (**Fig. 2B**). BOS also stimulated phagocytosis of bacteria by BMDM, THP-1 and HMDM **(Fig. S2A)**.

**Figure 2.**
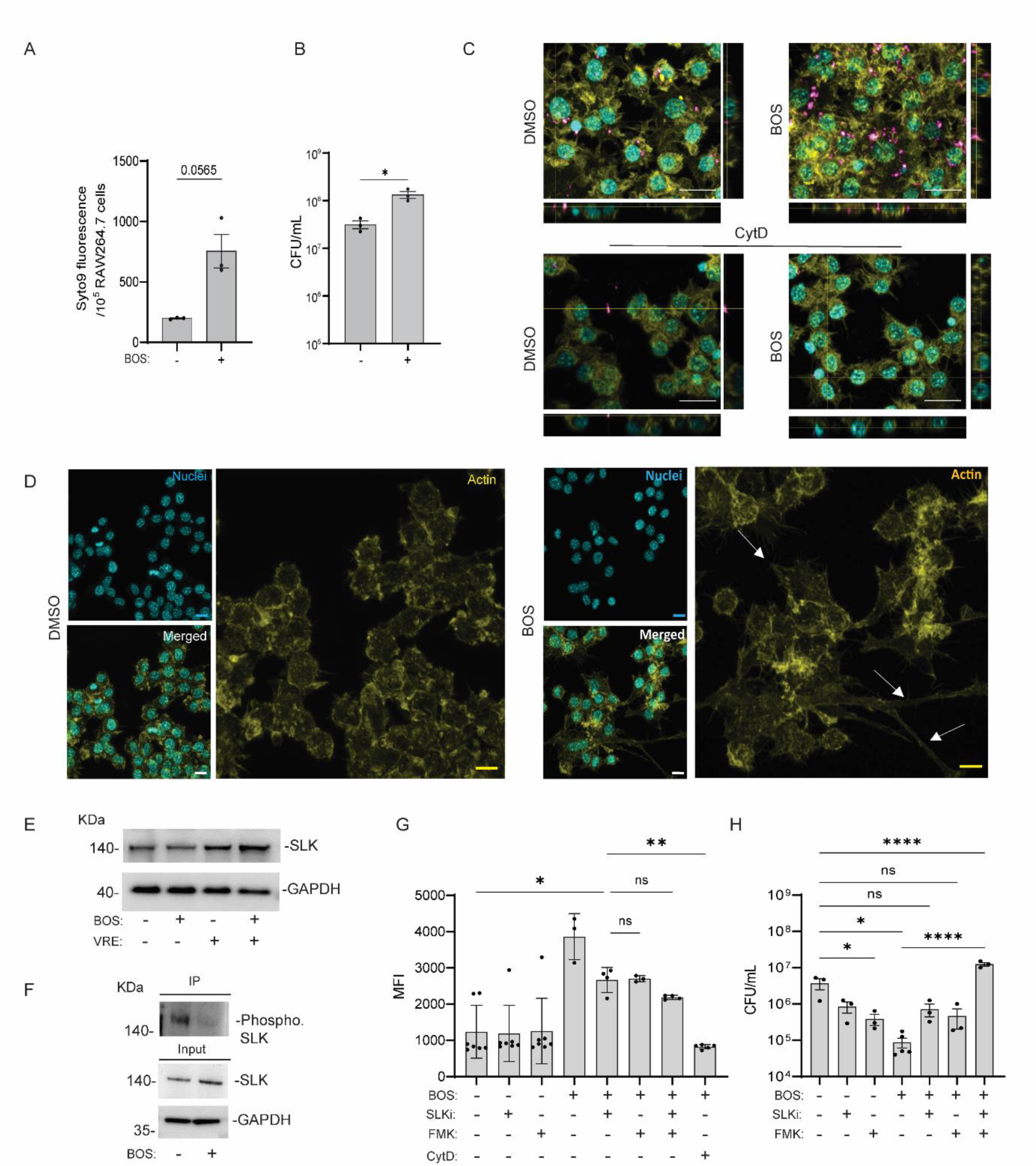
- BOS stimulates macrophage phagocytosis of bacteria through actin remodelling. **(A)** Comparison of uptake of SYTO9-labelled VRE by RAW264.7 macrophages with or without BOS pre-treatment. RAW264.7 macrophages with or without BOS pre-treatment were infected for 1 h with SYTO9-labelled VRE, followed by quenching extracellular fluorescence with trypan blue, and fluorescence intensity measurement by plate reader. **(B)** Comparison of VRE CFU after 1 h infection of RAW264.7 macrophages with or without BOS pre-treatment. Data (mean ± SEM) are summary of at least three independent experiments (A-B). **(C)** Representative CLSM images and orthogonal views of SYTO9-labelled VRE (pink) infected RAW264.7 macrophages with and without BOS pretreatment. CytD (40 μM) was added 30 min prior to infection. Samples were stained with phalloidin for actin visualization and Hoechst 33342 for DNA visualization. **(D)** Representative CLSM images of DMSO (left panels) or BOS (right panels) treated RAW264.7 macrophages that were stained with phalloidin (actin) and Hoechst 33342 (no infection). White arrows point to examples of cell projections. Images are maximum intensity projections of the optical sections (0.64 μm z-volume) and are representative of 3 independent experiments (C-D). Scale bar: 20 µm. **(E)** Western blotting analysis of SLK levels in whole-cell lysates. RAW264.7 cells with (+) and without (−) VRE infection were treated with BOS (+) or left untreated (−) and the lysates were subjected to Western blotting with anti–SLK and anti-GAPDH antibodies. **(F)** Immunoprecipitation of phosphorylated SLK in RAW264.7 cells following BOS treatment. RAW264.7 cells were treated with BOS (+) or left untreated (−), cell lysates were precipitated with anti-SLK antibody, followed by Western blotting with anti-phosphoserine/threonine antibody (top). Whole-cell lysates used for immunoprecipitation were subjected to Western blotting with anti–SLK and anti- GAPDH antibodies (bottom). **(G)** Inhibition of BOS-stimulated phagocytosis by various inhibitors. RAW264.7 macrophages with and without BOS pre-treatment were infected for 1h with SYTO9-labelled VRE in the presence or absence of various inhibitors. Samples were quenched with trypan blue followed by flow cytometry. Mean Fluorescence Intensity (MFI) is shown for samples that were not treated (DMEM) or pre-treated overnight with BOS (0.52 μg/mL), SLKi (1 μM), FMK (50 μM) alone or in combination. CytD (40 μM) was added 30min prior to VRE infection. **(H)** Inhibition of BOS-stimulated phagocytosis by various inhibitors. RAW264.7 cells were infected with VRE in the presence of BOS (0.52 μg/mL), SLKi (1 μM), FMK (50 μM) alone or in combination. Intracellular bacterial CFU was quantified after 18h. Data (mean ± SEM) are a summary of at least three independent experiments. Statistical analysis was performed using unpaired t test with Welch’s corrections (A-B), using ordinary one- way ANOVA, followed by Tukey’s multiple comparison test (G-H); NS, P > 0.05; *P ≤ 0.05, **P ≤ 0.01, and ****P ≤ 0.0001.

To further probe BOS-stimulated phagocytosis, we added the actin polymerization inhibitor Cytochalasin D (CytD) to RAW264.7 cells 30 min prior to infection with Syto9- stained VRE, and analyzed bacterial phagocytosis by microscopy and flow cytometry. CytD pre-treatment significantly inhibited phagocytosis of VRE by RAW264.7 cells (**Fig. 2C** **and** **S2B-C**). We also treated RAW264.7 macrophages with BOS or the vehicle control for 18 h, in the absence of infection, followed by phalloidin staining and confocal microscopy. BOS-treated RAW264.7 macrophages displayed spikier morphologies and long projections as compared to untreated cells (**Fig. 2D**). Similarly, BOS-treated HMDM cells also exhibited elongated morphology **(Fig. S2D)**, in agreement with our previous observation [26].

Studies have shown that BOS inhibits the phosphorylation of the SRC-CK2-SLK cascade [26]. Phosphorylation of SLK (also known as Ste20-like kinase) by CK2 inhibits SLK activity, SLK protein level, and, ultimately, its actin-remodelling activity [27]. To test whether inhibition of SLK phosphorylation is involved in BOS-induced actin-remodelling, we first determined whether BOS inhibits SLK phosphorylation. We precipitated SLK with an anti-SLK antibody, followed by Western blotting with anti-phosphoserine/threonine antibody. The level of SLK in RAW264.7 macrophages was similar in the presence or the absence of BOS and/or bacteria (**Fig. 2E**). However, with BOS treatment, SLK phosphorylation was greatly decreased (**Fig. 2F**).

SLK activity is also regulated by caspase 3 [28], which is known to be stimulated by BOS [29]. In BOS-treated RAW264.7 macrophages, we observed an increase in the level of activated (cleaved) caspase 3, but this increase was abolished when BOS- treated macrophages were infected **(Fig. S2E)**. Consistently, caspase 3 activity in cell lysates, measured by cleavage of the substrate DEVD-AFC (free AFC emits a yellow-green fluorescence), was significantly increased following BOS treatment, and was inhibited with the use of caspase-3 inhibitor Z-DEVD-FMK (FMK) **(Fig. S2F)**.

We further investigated the role of SLK and caspase 3 in phagocytosis and bacterial killing by macrophages using SLK and caspase 3 inhibitors SLKi and FMK, respectively. RAW264.7 macrophages were treated with BOS or vehicle alone plus or minus SLKi, FMK, or both for 18h, followed by addition of Syto9-stained VRE for 30min and flow cytometry. BOS-stimulated macrophage phagocytosis of VRE was partially inhibited by SLKi and FMK, more potently inhibited by both, and most dramatically inhibited by CytD (**Fig. 2G**). To assess intracellular bacterial killing, RAW264.7 macrophages were incubated with VRE for 3 h and then treated with BOS alone, BOS plus SLKi or plus FMK, or BOS plus both SLKi and FMK for 15 h, in the presence of gentamicin and penicillin, followed by quantification of intracellular CFU. BOS- stimulated phagocytosis and subsequent bacterial killing was partially inhibited by SLKi and FMK and completely inhibited by both inhibitors together compared to the untreated control (**Fig. 2H**). Neither inhibitor compromised macrophage viability as assessed by LDH assay **(Table S3)**. Together, these results show that BOS stimulates macrophage phagocytosis of bacteria by SLK-mediated actin remodelling.

### BOS induces macrophage expression of genes involved in bacterial uptake

To gain deeper insight into the mechanism by which BOS stimulates macrophage phagocytosis and subsequent killing of bacteria, we performed total RNA sequencing of DMSO control or BOS-treated RAW264.7 cells (18 h). Overall, transcription of 141 genes was upregulated and transcription of 135 genes was downregulated following BOS treatment **(Supplementary data 1)**. Among the differentially expressed genes (DEGs), pathways associated with cell-matrix adhesion, cell adhesion, and cell migration were upregulated (**Fig. 3A**), in agreement with our results of BOS on cell morphologies (**Fig. 2D**). Moreover, transcript levels of cell surface markers involved in bacterial uptake, killing, and presentation, including CD80, CD11b and TLR4, were significantly upregulated (by 1.06, 0.74, and 0.48-fold, respectively), while CD36 was downregulated by 2.50-fold **(Table S4)**. Consistently, flow cytometry staining confirmed the increase of CD11b and TLR4 and decrease of CD36 on the surface of BOS-treated macrophages **(Fig. S3A)**. In addition, several other markers involved in bacterial uptake and killing, including CD206, CD14, and Dectin-1, were increased following BOS treatment (**Fig. 3B** **and S3A)**. Similarly, CD14 and Dectin-1 levels were also upregulated on macrophages isolated from wounds of mice following a single IP injection of BOS (**Fig. 3C** **and** **S3B**). The overall percentage of macrophages and neutrophils in uninfected wounds was not affected by BOS treatment (**Fig. 3D**). Thus, although BOS does not stimulate recruitment of macrophages to the site of infection, it stimulates transcription of genes involved in bacterial uptake, further supporting a role for BOS in stimulating macrophage phagocytosis of bacteria.

**Figure 3.**
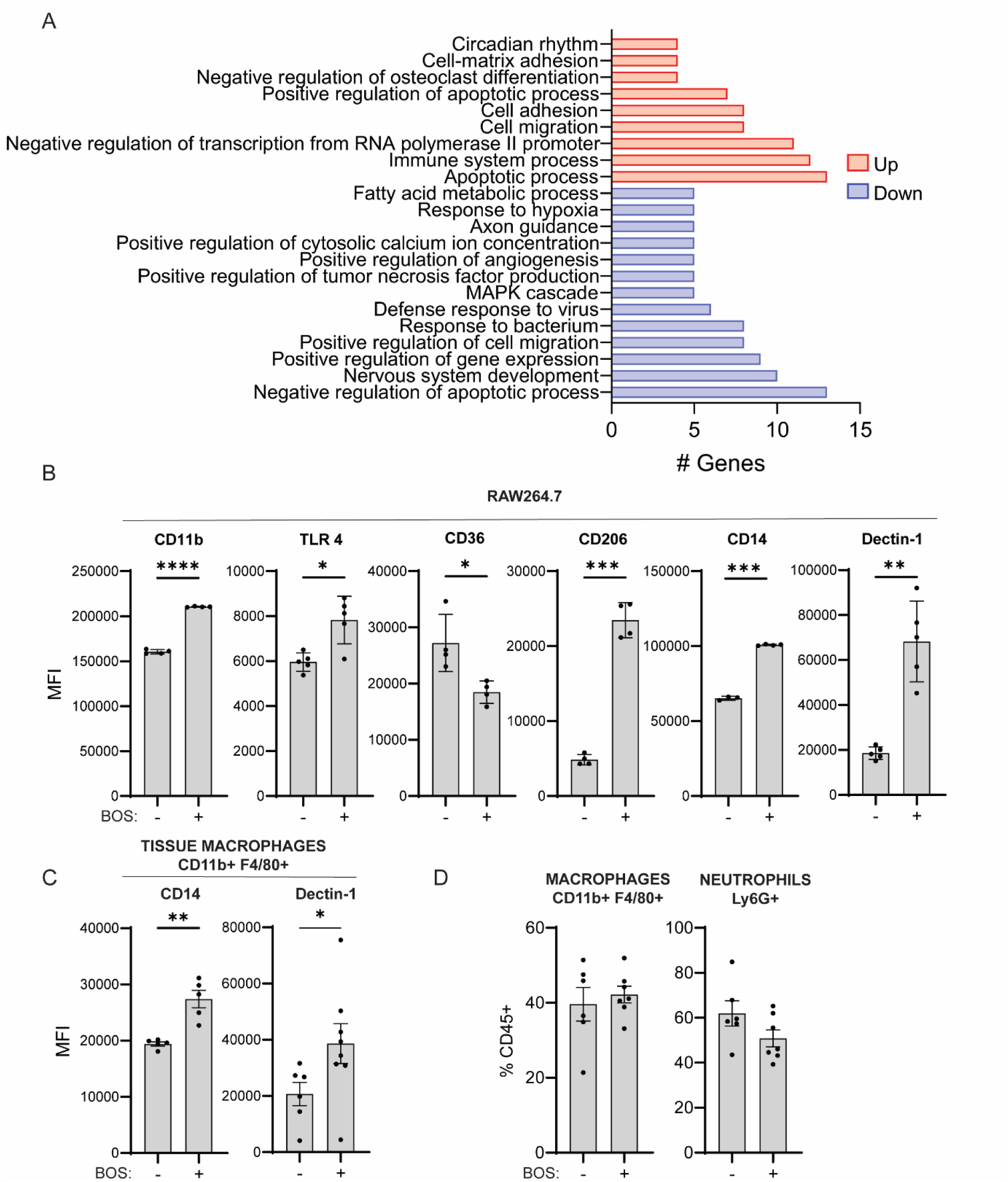
- BOS induces macrophage expression of genes involved in bacterial uptake. **(A)** Functional enrichment analysis of DEGs induced in RAW264.7 cells after 15 h treatment with BOS. **(B-C)** Comparison of mean fluorescence intensity (MFI) of CD11b, TLR4, CD36, CD206, CD14 and Dectin-1 staining gating on CD45^+^ RAW264.7 macrophages with or without BOS treatment (B) and CD14 and Dectin-1 on CD45^+^ CD11b^+^ F4/80^+^ macrophages from wounds of mice treated with an IP injection of vehicle (-) or BOS (+) (C). Data (mean ± SEM) are summary of at least two independent experiments (B-C) with two to four mice per experiment. **(D)** Relative levels of macrophages and neutrophils recovered from wounds of animals following IP injection with vehicle or BOS. Data (mean ± SEM) are a summary of at least two independent experiments. Each dot represents one mouse. Statistical analysis was performed using unpaired t test with Welch’s corrections. NS, P > 0.05; *P ≤ 0.05, **P ≤ 0.01 and ***P ≤ 0.001.

### BOS stimulates macrophage killing of bacteria via reactive oxygen species

Reactive oxygen species (ROS) and lysosomal activity are two crucial mechanisms of intracellular bacterial killing by macrophages [7]. Following BOS treatment of RAW264.7 cells, the phagolysosomal proteins LAMP-1, cathepsin B (CtsB), CtsD, Rab5, Rab7 were unchanged, whereas levels of rubicon, a protein involved in non-canonical phagocytosis, was elevated **(Fig. S4A)** [30]. It was previously reported that BOS can induce leakage of lysosomal enzymes into cytosol [29]. However, addition of CtsD and B inhibitors pepstatin A (Peps) and CA-074 into BOS-treated macrophages did not affect the killing of intracellular bacteria **(Fig. S4B)**.

To investigate the role of ROS in BOS-stimulated bacterial killing by macrophages, we first tested if BOS induces ROS in macrophages, independent of infection. RAW264.7 macrophages were treated with BOS for 18 h, followed by quantification of DHR123 fluorescence. BOS stimulated ROS production, which was reduced by N-acetyl cysteine (NAC) (**Fig. 4A**). Similarly, when ROS was measured by DCFDA fluorescence, BOS also stimulated ROS production although the increase did not reach significance (**Fig. 4B**). However, when BOS-treated RAW264.7 macrophages were infected with VRE for 3 h, ROS level was significantly elevated, reaching the level induced by TBHP. Live cell imaging using CellRox to label ROS, LysoTracker to label lysosomes, CellTracker to label cells, and GFP-expressing VRE, supported that BOS stimulates production of ROS, which often co-localized with lysosome **(Fig. S4C)**. In the presence of infection and BOS treatment, we observed more ROS, which was co-localized with both lysosomes and bacteria. Consistently, when ROS was quenched by either NAC or TEMPO, BOS-stimulated killing of bacteria by macrophages was abolished (**Fig. 4B**-**C**). Thus, BOS stimulates macrophage killing of phagocytosed bacteria by ROS.

**Figure 4.**
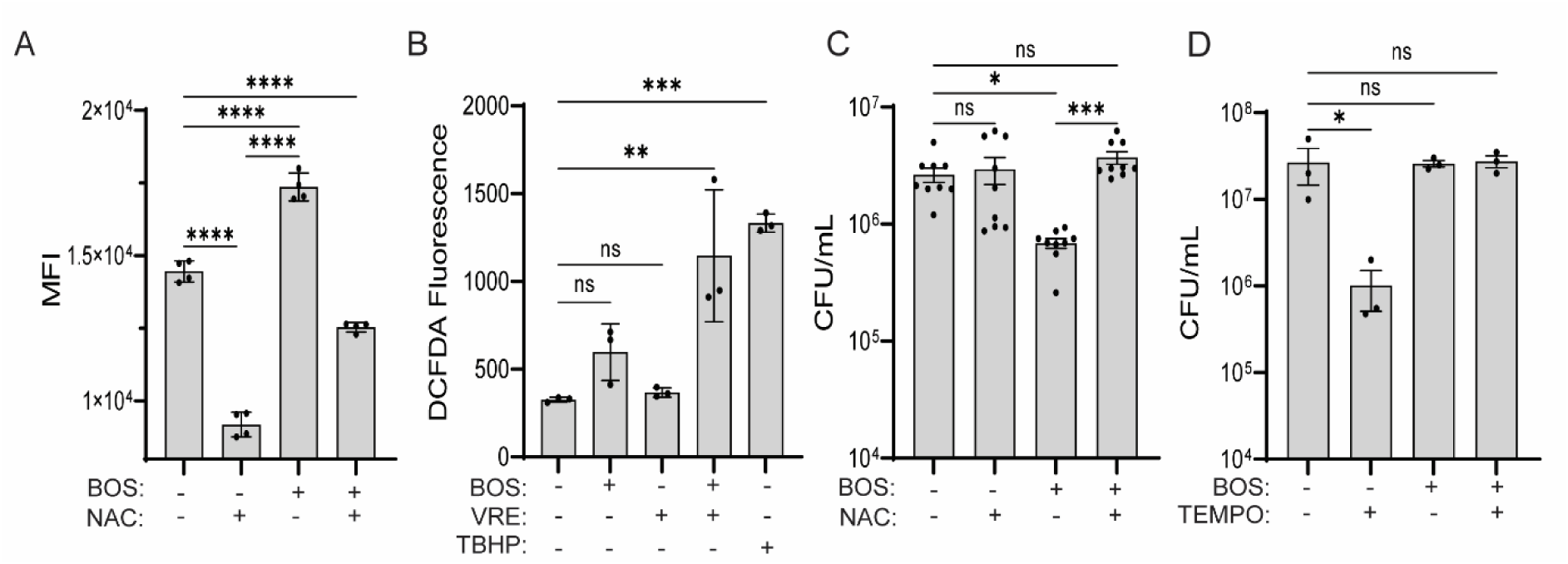
- BOS stimulates macrophage killing of bacteria via ROS. **(A)** ROS levels as measured by flow cytometry of DHR123 fluorescence. RAW264.7 macrophages were treated with BOS alone or in combination with NAC (5 mM) overnight, followed by flow cytometry. **(B)** ROS levels as measured by plate reader of DCFAD fluorescence. RAW264.7 macrophages were left untreated or treated with BOS or tert-butyl hydroperoxide (TBHP, 100 μM, positive control), with or without VRE infection for 3 h. **(C-D)** BOS-stimulated bacterial killing by macrophages is abolished by neutralization of ROS. RAW264.7 cells were infected with VRE in the presence of BOS (0.52 μg/mL), NAC (5 nM), or TEMPO (50 μM), alone or in combination. Intracellular bacterial CFU was quantified after 15 h. Data (mean ± SEM) are a summary of at least three independent experiments. Statistical analysis was performed using ordinary one-way ANOVA, followed by Tukey’s multiple comparison test; NS, P > 0.05; *P ≤ 0.05, **P ≤ 0.01, and ***P ≤ 0.001.

### BOS promotes survival of infected macrophages

To investigate the effect of infection and BOS treatment on macrophages, we also performed RNA sequencing of VRE-infected RAW264.7 macrophages following 15 h of BOS exposure, or DMSO control. Transcription of 40 genes was upregulated and transcription of 99 genes was downregulated **(Supplementary data 2)**. Among the DEGs, pathways associated with microtubule-based processes and cytoskeleton organization were upregulated (**Fig. 5A**), consistent with BOS-induced actin remodelling. Pathways associated with stress responses (response to bacterium, response to oxygen-containing compound, response to organic substance, and cellular response to chemical stimulus) and protein metabolism (regulation of cellular protein metabolic process and regulation of protein metabolic process) were also upregulated (**Fig. 5A**), suggesting that BOS may affect survival of infected macrophages.

**Figure 5.**
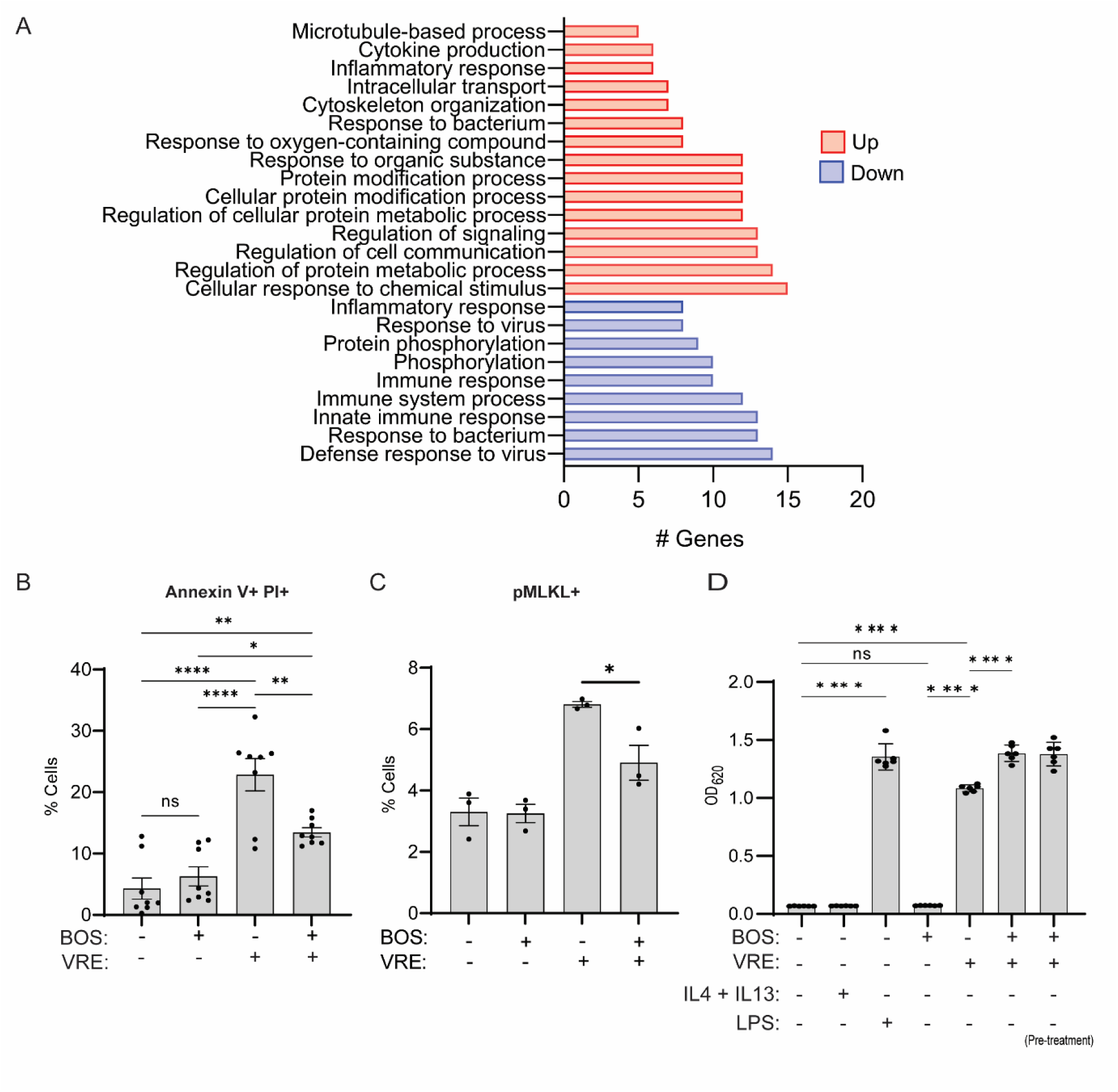
- BOS promotes survival of infected macrophages. **(A)** Functional enrichment analysis of DEGs induced by BOS treatment of VRE-infected RAW264.7 cells. **(B-C)** Comparison of percentages of Annexin V^+^ and PI^+^ (B) and pMLKL^+^ (C) cells. RAW264.7 cells were not infected or infected with VRE in the presence or absence of BOS. Cell viability was assayed by Annexin V and PI staining and expression of pMLKL was assayed by intracellular staining followed by flow cytometry. **(D)** NF-κB activation measurement. RAW267.4 macrophages were untreated or treated with BOS or LPS (100 ng/mL) or IL-4 (10 ng/mL) and IL-13 (10 ng/mL) for 18h prior to measurement of NF-κB-driven SEAP reporter activity. When the effect of VRE infection in RAW234.7 cells was evaluated, NF-κB-driven SEAP reporter activity was measured at the end of the intracellular infection assay with BOS treatment performed at the time of infection or prior to the start of the experiment (pre-treatment). Data (mean ± SEM) are a summary of at least three independent experiments. Statistical analysis was performed using ordinary one-way ANOVA, followed by Tukey’s multiple comparison test; NS, P > 0.05; *P ≤ 0.05, **P ≤ 0.01, ***P ≤ 0.001 and ****P ≤ 0.0001.

To test this hypothesis, we monitored induction of apoptosis and membrane permeability as an indication of cell viability in infected macrophages in the presence or absence of BOS. While BOS did not affect RAW264.7 viability in the absence of infection (**Fig. 5B**), Annexin V^+^ PI^+^ cells were significantly increased following infection. BOS treatment of infected cells reduced the percentage of Annexin V^+^ PI^+^ cells by ∼50%, consistent with delayed apoptosis and prolonged viability. Similarly, BOS exposure alone did not affect the level of phosphorylated MLKL (pMLKL), a marker of necroptosis in macrophages [31] (**Fig. 5C** **and** **S5A**), VRE infection significantly induced the level of pMLKL, and BOS treatment of infected cells reduced pMLKL by ∼30%.

Pathways associated with cytokine production and inflammatory response were also upregulated in VRE-infected RAW264.7 macrophages that were treated with BOS (**Fig. 5A**). Consistently, induction of NF-κB activity as measured by reporter assay was induced by VRE infection and further enhanced by BOS treatment (**Fig. 5D**). NF-κB activation can promote cell survival [32]; therefore, we measured the effect of an NF-κB inhibitor, the quinazoline derivative compound QNZ, on the survival of infected macrophages with or without BOS treatment. 18h after VRE infection, the percentage of Annexin V^+^ PI^+^ RAW264.7 macrophages was ∼45%, which was reduced to 15% in the presence of BOS **(Fig. S5B)**. In the presence of QNZ, the percentage of Annexin V^+^ PI^+^ of BOS-treated infected macrophages was increased to ∼35%. Furthermore, QNZ did not inhibit macrophage killing of VRE in the absence of BOS, but inhibited BOS-stimulated macrophage killing of bacteria **(Fig. S5C)**. Together, these results show that BOS promotes survival of infected macrophages through NF-kB regulated pathways.

### Other Src kinase family inhibitors also stimulate macrophage killing of bacteria

Our findings thus far suggest that inhibition of the SRC kinase in macrophages improved bacterial uptake and elimination. Therefore, we tested whether other Src family kinase inhibitors also promote macrophage killing of bacteria. RAW264.7 macrophages were infected with VRE for 3 h and then treated with DMAT (a CK2 inhibitor), saracatinib (SARA), dasatinib (DASA) and tirbanibulin (TIR), all inhibitors of SRC. Although all four compounds stimulated RAW264.7 killing of bacteria, only DMAT-stimulated killing reached statistical significance (**Fig. 6A**). However, both DMAT and TIR stimulated significantly more killing of bacteria by HMDM *in vitro* (**Fig. 6B**). Similarly, a single IP dose (5mg/kg) of either DMAT or TIR reduced VRE CFU of wound infection in mice by 1.5-log and 1-log, respectively (**Fig. 6C**). At the dosage used, these compounds did not cause cell death as measured by the LDH assay **(Table S4)**. All four compounds, except DMAT, significantly stimulated phagocytosis of bacteria by RAW 264.7 cells (**Fig. 6D**). Furthermore, all compounds stimulated the expression of Dectin-1 (**Fig. 6E**), and DMAT, DASA and TIR also stimulated CD14 expression (**Fig. 6F**). Thus, other inhibitors of the Src family kinases also stimulate macrophage killing of bacteria *in vitro* and *in vivo*.

**Figure 6.**
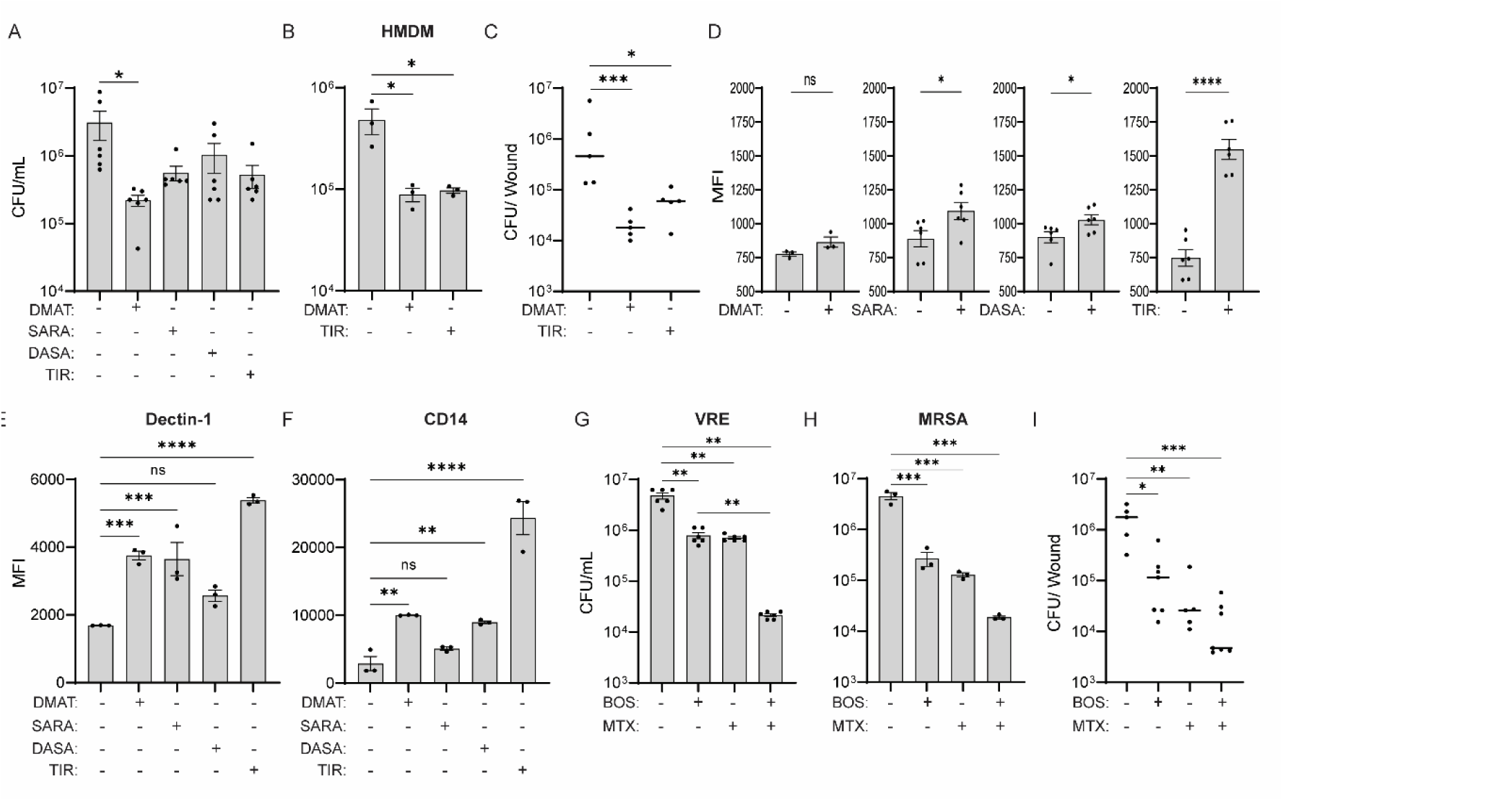
- Other SFK inhibitors also stimulate macrophage killing of bacteria. **(A)** Comparison of VRE CFU in RAW264.7 cells untreated or treated with various Src kinase inhibitors. Src kinase inhibitors used were: DMAT (1 μM), SARA (1 μM), DASA (1 μM) or TIR (0.33 μM). Data (mean ± SEM) are a summary of at least three independent experiments. **(B)** Comparison of VRE CFU in HMDM that were treated with vehicle, DMAT (1 μM) or TIR (0.33 μM). **(C)** Comparison of VRE CFU per infected wound of animals treated with an IP injection of DMSO, DMAT (5 mg/kg) or TIR (5 mg/kg). Data were from two independent experiments with two to three mice per experiment. Each symbol represents one mouse, with the median indicated by the horizontal line. **(D)** Phagocytosis of SYTO9-labelled VRE by RAW264.7 macrophages in the presence or absence of various inhibitors. Data (mean ± SEM) are a summary of at least three independent experiments. **(E-F)** Comparison of MFI of CD14 and Dectin-1 staining of CD45^+^ RAW264.7 macrophages non-treated or treated with Src kinase inhibitors. Data (mean ± SEM) are a summary of at least three independent experiments. Statistical analysis was performed using ordinary one-way ANOVA, followed by Tukey’s multiple comparison test (A, E, F), or using Kruskal-Wallis test with uncorrected Dunn’s posttest (C) or using unpaired t test with Welch’s corrections (D); NS, P > 0.05; *P ≤ 0.05, **P ≤ 0.01, and ****P ≤ 0.0001. **(G-H)** Comparison of VRE (E) and MRSA (F) CFU in RAW264.7 that were treated with BOS or MTX (0.515 μg/ml) or in combination. **(I)** Comparison of VRE CFU per infected wound of animals treated with five IP injections of vehicle (DMSO) or BOS alone or in combination with five topical treatments of MTX (0.515 μg/ml). Data were from two independent experiments with two to three mice per experiment. Each symbol represents one mouse, with the median indicated by the horizontal line. Statistical analysis was performed using ordinary one-way ANOVA, followed by Tukey’s multiple comparison test (G-H) or using Kruskal-Wallis test with uncorrected Dunn’s posttest (I). NS, P > 0.05; *P ≤ 0.05, **P ≤ 0.01, and ****P ≤ 0.0001.

### BOS and mitoxantrone are additive in promoting macrophage clearance of bacteria

We previously reported that mitoxantrone (MTX) stimulates macrophage killing of bacteria by inducing the expression of lysosomal enzymes without inducing phagocytosis [1]. Since BOS stimulates macrophage killing of bacteria by stimulating phagocytosis without inducing expression of lysosomal enzymes, we tested whether the two compounds are synergistic or additive in stimulating macrophage killing of bacteria. RAW264.7 macrophages were infected with VRE or MRSA for 3 h and then treated with BOS alone, or MTX alone, or both for 15h, followed by intracellular CFU enumeration. BOS or MTX reduced VRE CFU by about 0.8-log whereas the two compounds together reduced CFU by 2.3-log (**Fig. 6G**). Similarly, BOS or MTX reduced MRSA CFU by about 1.2-log whereas the two compounds together reduced CFU by 2.3-log (**Fig. 6H**). In the mouse model of wound infection with VRE, five treatments with BOS alone (IP) or MTX alone (topical) reduced bacterial burden in the wounds by 1.2-log and 1.6-log, respectively (**Fig. 6I**). Combination treatment with the same dosing regimen reduced VRE CFU in wounds by 2.1-log. These results show that BOS and MTX are additive and can be used in combination to improve bacterial clearance *in vitro* and *in vivo*.

## Discussion

Widespread antimicrobial resistance is a major global health challenge. Compounds that enhance host immune responses are emerging as alternative approaches to treat antibiotic resistant infections. BOS is a chemotherapeutic for treating adults with Philadelphia chromosome-positive (Ph+) chronic myelogenous leukemia through its inhibition of the SRC/ABL tyrosine kinases [33]. In this study, we show that BOS has potential as a host-targeted therapy for the control of bacterial infection since it does not possess antibiotic activity but enhances macrophage phagocytosis and intracellular killing of bacteria, as well as the survival of infected macrophages.

We show that BOS stimulates macrophage clearance of bacterial infection both *in vitro* and *in vivo*. BOS stimulated the effective clearance of VRE, MRSA, *P. aeruginosa* and the multi-drug resistant *E. coli* strain 958 by the murine macrophage cell line RAW264.7 *in vitro*. Similarly, BOS stimulated a significantly more effective clearance of VRE by murine BMDM, the human monocytic cell line THP-1 and HMDM *in vitro*. Pre-treatment of RAW264.7 and BMDM with BOS overnight was sufficient to enhance bacterial killing. Furthermore, mice infected with VRE and treated with a single intraperitoneal dose of BOS showed a significant reduction in bacterial load 24 h later. A similar efficacious effect was also observed with multiple low-dose topical applications on the infected wounds. Importantly, depletion of macrophages in the wounds confirmed that these cells are required for BOS-stimulated bacterial clearance.

We show that BOS stimulates bacterial clearance through three mechanisms. First, BOS promotes macrophage phagocytosis of bacteria by inducing actin remodelling and upregulating bacterial uptake markers. Studies have shown that BOS inhibits SRC phosphorylation, affects actin remodelling, and affects cell morphology [26, 34]. Past reports also showed that phosphorylation of SLK is dependent on the SRC-CK2-SLK signalling pathway [27]. Moreover, SLK phosphorylation, together with caspase 3 activity, directly impacts SLK function [27, 28]. Consistent with these previous studies, we showed that BOS treatment inhibits SLK phosphorylation, leading to extensive actin remodeling, and increased phagocytosis activity, which were abolished by the actin polymerization inhibitor Cytochalasin D and partially inhibited by SLK and caspase 3 inhibitors. With this study, we provide enough evidence to fully bridge the SRC-CK2-SLK phosphorylation cascade to actin remodelling and enhanced phagocytic activity. Furthermore, we show that BOS treatment upregulates expression of cell surface markers involved in bacterial uptake, killing, and presentation, including CD80, CD11b, TLR4, CD206, CD14 and Dectin-1 both *in vitro* and *in vivo*. Together, the increased recognition and uptake of bacteria, together with increased actin-remodeling, likely result in the increased phagocytosis of bacteria by BOS-treated macrophages, leading to more effective clearance of bacteria.

Second, BOS stimulates macrophage killing of phagocytosed bacteria via increased production of reactive oxygen species. BOS stimulated macrophage ROS production in the absence of infection and even more dramatically in the presence of infection. Live cell imaging showed that ROS colocalized with bacteria in the lysosomes. When ROS was quenched using NAC or TEMPO, intracellular bacterial killing was also reduced, demonstrating a significant role of ROS in enhancing the bactericidal activity of BOS-treated macrophages.

Third, BOS stimulates survival of infected macrophages, therefore clearance of infection. Transcriptional profiling of infected BOS-treated macrophages showed that pathways associated with stress responses and protein metabolism were upregulated. Consistently, we observed reduced cell death of infected macrophages following BOS treatment. As NF-κB activation is known to promote cell survival [32], we directly showed induction of NF-κB activity by BOS using a reporter assay. Furthermore, inhibition of NF-κB activity by QNZ partially blocked the effect of BOS in promoting the survival of infected macrophages and killing of bacteria. Together, these three mechanisms contribute to the observed effect of BOS on promoting macrophage clearance of bacterial infection.

We also observed enhanced phagocytic activity and intracellular bacterial killing by other Src kinase inhibitors. DMAT inhibits the kinase CK2. Although CK2 is not strictly considered part of the Src kinase family, it is a close partner downstream of different pathways coordinated by Src family members [35]. DMAT stimulated clearance of VRE by RAW264.7 cells *in vitro* and in a murine wound infection model. SARA, DASA and TIR also inhibit SFKs. SARA is the most promiscuous of these inhibitors and can inhibit several members of SFKs [36]. DASA and TIR are more specific to SRC [37, 38]. We showed that some of these inhibitors significantly stimulated phagocytosis of bacteria by RAW 264.7 cells and the expression of Dectin-1. DMAT and TIR also stimulated significantly more killing of bacteria by HMDM *in vitro* and a single IP dose significantly reduced VRE burden in wound infection in mice. Interestingly, a previous study showed that lower doses of DASA can help in a sepsis model of polymicrobial infection and can enhance the phagocytic activity of neutrophils [18]. Since SRC-mediated pathways can collaborate or interact with other signalling pathways, the impact of inhibiting SRC likely varies in different cell types based on the level and activity of SRC partners and the level of inhibition that can be achieved [8]. Nevertheless, the similar effects of other Src kinase inhibitors provide further support for a critical role of the phosphorylation cascade axis SRC-CK2-SLK in mediating the observed effect of BOS.

Finally, we show that BOS and mitoxantrone (MTX) exhibit an additive effect in promoting macrophage clearance of bacterial infection both *in vitro* and *in vivo.* We have previously shown that MTX, a chemotherapeutic used to treat advanced prostate cancer and acute nonlymphocytic leukemia, has antibiotic activity as well as stimulates macrophage recruitment to the site of infection and killing of bacteria by inducing the expression of lysosomal enzymes [1]. In contrast, BOS stimulates phagocytosis, intracellular ROS production and survival of infected macrophages. The observed additive effect between BOS and MTX likely results from their non-overlapping effects on macrophages. These findings open possibilities of reducing bacterial drug resistance by combination use where BOS complements the action of other compounds and antibiotics. As an adjuvant, BOS stimulates host macrophage clearance of bacteria and therefore provides a valuable addition for therapies that aim to target the bacteria in order to achieve complete elimination of infection [39]. Thus, our study of BOS adds a new tool to combat bacterial drug-resistant by boosting host innate immunity.

## Materials and Methods

### Bacterial strains and growth conditions

Bacterial strains used in this study are listed in **Table S1**. Bacterial strains were grown using Brain Heart Infusion (BHI) broth and agar (Becton, Dickinson and Company). Bacterial strains were streaked from glycerol stocks stored at -80 °C, inoculated and grown overnight statically for 16-20 h either in 10 mL of liquid BHI broth or DMEM (Gibco, high glucose and no sodium pyruvate) + 10% FBS medium. Cells were harvested by centrifugation at 8000 RPM (25°C) for 5 min. The supernatant was discarded, and the pellet was then resuspended in either DMEM + 10% FBS or sterile PBS to an optical density at 600 nm (OD_600nm_) of 0.7 for VRE, equivalent to 2–3×10^8^colony forming units (CFU).

### Antimicrobial and minimum inhibitory concentration assays

Bacterial growth assays were carried out in complete DMEM medium as described previously [40]. 2 μl of overnight cultures grown in DMEM were added to 200 μl of medium in a 96-well plate with increasing concentrations of BOS and/or antibiotics. The OD_600_ at the zero-time point was established. Bacteria were grown statically 96- well plates at 37°C for up to 24 h. Final OD_600_ measurements were acquired using a Tecan M200 microplate reader.

### Human monocyte derived macrophages (HMDM) and cell lines

Isolated peripheral blood (PB) primary human monocytes were purchased from StemCells Technologies. For *in vitro* differentiation of monocytes into human macrophages, isolated monocytes were cultured in complete RPMI1640 supplemented with 10% heat-inactivated fetal bovine serum (FBS) (PAA, GE Healthcare), 2 mM L-glutamine (Corning) and 1% PenStrep solution (Gibco) in the presence of 50 ng/mL recombinant human M-CSF (Biolegend) for 7 days. The RAW264.7 murine macrophage-like cell line (InvivoGen), and the THP-1 monocytic cells derived from an acute monocytic leukemia patient cell line (ATCC), were cultured at 37°C in a 5% CO_2_ humidified atmosphere. THP-1 cells were cultured in complete RPMI1640 supplemented with 10% heat-inactivated fetal bovine serum (FBS). THP-1 cells were differentiated to macrophages with 100 ng/mL phorbol-12-myristate 13- acetate (PMA, Sigma-Aldrich) for 3 days. RAW264.7 cells were grown and maintained in Dulbecco’s modified Eagle’s medium (DMEM) (Gibco, high glucose, no sodium pyruvate) with 10% heat-inactivated FBS (PAA, GE Healthcare), and 100 U of penicillin–streptomycin (Gibco, Thermo Fisher Scientific). The culture medium was replaced every three days, and upon reaching 80% confluency, cultures were passaged. RAW264.7 cells passaging was achieved by gentle cell scraping and seeding cells at a density of 3×10^6^ cells/T75 flask (Nunc; Thermo Fisher Scientific).

### Mouse bone marrow-derived macrophages (BMDM)

BMDMs were prepared as described previously [41]. Briefly, fresh bone marrow cells were isolated from mice, plated in complete RPMI with 50 ng/mL recombinant M-CSF (Biolegend) and cultured for 6 days with medium change every 3 days.

### Intracellular infection assay

Intracellular infection assays were performed as described in [22] with some modifications. Cells were seeded at a density of 10^6^ cells/well or 8×10^5^ cells/well in a 6-well or 96-well tissue culture plate (Nunc; Thermo Fisher Scientific), respectively, and allowed to attach overnight at 37 °C in a 5 % CO_2_ humidified atmosphere. Cells were infected at a multiplicity of infection (MOI) of 10 for up to 3 h. Following infection, the media was aspirated, and the cells were washed three times in PBS and incubated with 150 μg/ml of gentamicin (Sigma-Aldrich) and 50 μg/ml penicillin G (Sigma-Aldrich) to kill extracellular bacteria and BOS (0.52 μg/ml) in complete DMEM for 15-18 h to selectively kill extracellular bacteria. The antibiotic-containing medium was then removed, and the cells were washed 3 times in PBS before addition of 2% Triton X– 100 (Sigma-Aldrich) PBS solution to lyse the cells for enumeration of the intracellular bacteria. Variations of this assay included pre-treatment of mammalian cells, prior to bacterial infection, with BOS (0.52 μg/ml) followed by antibiotic treatment only, or co-treatment of cells at the time of infection with 50mM of NAC (Sigma) and the compounds listed in Table S4.

### Ethics statement

All animal experiments were performed with approval from the Institutional Animal Care and Use Committee (IACUC) in Nanyang Technological University, School of Biological Sciences under protocol ARF-SBS/NIE-A19061.

### Mouse wound excisional model

The procedure for mouse wound infections was modified from a previous study [42]. Briefly, male C57BL/6 mice (6-8 weeks old, 22 to 25 g; NTU, Singapore) were anesthetized with 3% isoflurane. Following dorsal hair trimming, the skin was then disinfected with 70% ethanol before creating a 6-mm full-thickness wound using a biopsy punch (Integra Miltex). Bacteria (1 x 10^6^ CFU) were added to the wound site. Then, the wound site was sealed with a transparent dressing (Tegaderm 3M). The IP injections of 30 μL of DMSO (vehicle), 30 μL BOS (5mg/kg), 30 μL DMAT (5mg/kg) or 30 μL TIR (5mg/kg) were performed prior to punching the wound. When preventive treatment was tested, DMSO or BOS were injected 24 h before infection, and when co-treatment with MTX was performed. When multiple treatments with BOS were performed, an 8mm Finn Chamber on Scanpor was placed around the wound to facilitate the removal of the transparent dressing for each treatment without disruption of the underlying bacterial biofilm. In total, five daily treatments of BOS IP injections (5mg/kg) were performed. Alternatively, five daily topical applications of 10 μL of PBS or 10 μL BOS (0.52 μg/mL) and/or MTX (0.515 μg/mL) were applied on the wound. After 24h or 4 days post-infection, mice were euthanized and a 1 cm by 1 cm squared piece of skin surrounding the wound site was excised and collected in sterile PBS. Skin samples were homogenized, and the viable bacteria enumerated by plating onto BHI plates.

### Phagocytosis Assay

*E. faecalis* V583 cells were fixed with 4% PFA for 15 min and washed thrice with PBS, prior to labelling with the membrane-permeant DNA dye - Syto9 (Thermo Fisher Scientific). Bacterial cells were then washed thrice with PBS and resuspended in DMEM + 10% FBS. Cells were infected with MOI10 of Syto9-labelled bacterial cells and incubated for 30 min to 1 hour at 37°C and 5% CO_2_. Following supernatant removal, infected cells were harvested and resuspended in PBS. The fluorescence of bacteria either free in the medium or attached to host cell membranes were quenched with a final concentration of 0.01 % trypan blue. As trypan blue cannot enter viable eukaryotic cells, the unquenched fluorescence reflected the bacterial cells that were internalized in viable cells. After staining, cells were immediately run through the flow cytometer. All data were collected using the BD LSRFortessa X-20 Cell Analyzer and analyzed with FlowJo V10.8.1 (BD Biosciences, USA). The samples were initially gated side scatter area (SSC-A) by forward scatter area (FSC-A) to select the macrophage populations. The cells’ population was subsequently gated forward scatter width (FSC-W) by side scatter area (SSC-A) to remove doublet populations. The resulting singlet cell population was then assessed for Syto9 fluorescent marker.

### Western Immunoblotting

Whole cell (WC) lysates were prepared by adding 488 μl of RIPA buffer (50 mM Tris-HCl, pH 8.0; 1 % Triton X–100; 0.5 % Sodium deoxycholate; 0.1 % SDS; 150 mM NaCl) to the wells after intracellular infection assays, where cells were scraped and kept in RIPA buffer for 30 min at 4 °C. Prior to the addition of 74.5 μl of 1 M DTT and 187.5 μl NuPAGE LDS Sample Buffer (4X) (Thermo Fisher Scientific), cells were further mechanically disrupted by passing the lysate through a 26g size needle. Samples were then heated to 95°C for 5 min. 15 μl of cell lysate proteins were then separated in a 4–12% (w/v) NuPAGE Bis-Tris protein gel and transferred to PVDF membranes. Membranes were incubated with Tris-buffered saline, TBS (50 mM Tris, 150 mM NaCl, pH 7.5) containing 0.1% (v/v) Tween-20 (TBST) and 5% (w/v) BSA for 1 h at room temperature. Membranes were incubated with 1:1000 primary antibodies in TBST containing 1% (w/v) BSA overnight at 4°C. Membranes were washed for 60 min with TBST at room temperature and then incubated for 2h at room temperature with goat anti-rabbit (H+L) HRP-linked secondary antibody (Invitrogen) or goat anti-mouse HRP-linked secondary antibodies (Invitrogen). After incubation, membranes were washed with TBST for 30 min and specific protein bands were detected by chemiluminescence using SuperSignal West Femto maximum sensitivity substrate (Thermo Fisher Scientific. All primary antibodies used in this study were monoclonal antibodies raised in rabbits except for the monoclonal antibody anti-phosphoserine/threonine which was raised in mice. All primary antibodies but anti- Nox2 (Invitrogen), polyclonal anti-SLK (Thermo Fisher Scientific) and anti-phosphoserine/threonine (Thermo Fisher Scientific) were purchased from Cell Signaling Technology.

### Immunoprecipitation of SLK from RAW264.7 cells

Samples were immuno-precipitated with Protein A/G Mag Sepharose (Abcam) and polyclonal anti-SLK (Thermo Fisher Scientific) according to the beads’ manufacturer’s instructions. Following incubation of 20 μl of magnetic bead slurry with 5 μg of anti-SLK for 1h at room temperature, 500 μl of whole cell lysates of BOS-treated or non-treated RAW264.7 cells in RIPA buffer was added and incubated overnight at 4°C. Using a magnetic particle concentrator, beads were washed twice with PBS and SLK recovered following the addition of 100 μl 0.1M glycine-HCl (pH 2.5 to 3.1). Samples were then subjected to western blot for detection of phosphoserine/threonine.

### Caspase-3 activity assay

Caspase 3 activity was measured using the Caspase-3 Activity Assay Kit (Fluorometric) (Abcam) as per the manufacturer’s instructions. Cells were treated with BOS overnight prior to caspase 3 activity measurement. FMK caspase inhibitor (50 mM) was added as negative control.

### Fluorescence Staining

Cells were seeded at 2×10^5^ cells/well in a 24-well plate with 10 mm coverslips, and allowed to attach overnight at 37°C and 5 % CO2. Vehicle (DMSO) or BOS treatment was performed overnight prior to fixation with 4% PFA at 4°C for 15 min. Cells were then blocked with PBS supplemented with 0.1% saponin and 2% bovine serum albumin (BSA). For actin labelling, the phalloidin–Alexa Fluor 555 conjugate (Thermo Fisher Scientific) was diluted 1:40 in blocking solution and incubated for 1h. Coverslips were then washed 3 times in PBS with 0.1 % saponin. For nuclei staining, Hoechst 33342 was added in a dilution of 1:1000. Coverslips were then subjected to a final wash with PBS, thrice. Finally, the coverslips were mounted with SlowFade Diamond Antifade (Thermo Fisher Scientific) and sealed. When bacteria were visualised, infection with VRE expressing episomally encoded GFP (pDasherGFP) [43] was performed with MOI10 for 1 h. Similarly, when ROS and lysosomes were visualized using CellRox at final concentration of 5μM (Invitrogen) and LysoTracker at a final concentration of 50 nM (Invitrogen), an hour incubation was performed before live imaging of these cells, which were not mounted. Celltracker was added at a final concentration of 2.5 μM, the night before ROS and lysosomes were visualised. Confocal images were then acquired on a 63x/NA1.4, Plan Apochromat oil objective fitted onto an Elyra PS.1 with LSM 780 confocal unit (Carl Zeiss), using the Zeiss Zen Black 2012 FP2 software suite. Laser power and gain were kept constant between experiments. Z-stacked images were processed using Zen 2.1 (Carl Zeiss). Acquired images were visually analyzed using ImageJ [44].

### Flow cytometry

Flow cytometry was performed as described in [22] with some modifications. Excised skin samples were placed in 1.5 ml Eppendorf tubes containing 2.5 U/ml liberase prepared in DMEM with 500 μg/ml of gentamicin and penicillin G (Sigma-Aldrich). The mixture was then transferred into 6-well plates and incubated for 1 h at 37°C in a 5% CO_2_ humidified atmosphere with constant agitation. Dissociated cells were then passed through a 70 μm cell strainer to remove undigested tissues and spun down at 1350 RPM for 5 min at 4°C. The enzymatic solution was then aspirated, and cells were blocked in 500 μl of FACS buffer (2% FBS and 0.2 mM ethylenediaminetetraacetic acid (EDTA) in PBS (Gibco, Thermo Fisher Scientific)). 10^7^ cells per sample were then incubated with 10 μl of Fc-blocker (anti-CD16/CD32 antibody, Biolegend) for 30 min, followed by incubation with an anti-mouse CD45, CD11b, and Ly6G (neutrophils), or CD45, CD11b and F4/80 (macrophages) plus CD14, Dectin-1, CD80, CD36, TIM-4, TLR-4, TLR-2, CD86, MHCII, CD163, or CD206 markers conjugated antibodies (Biolegend) (1:100 dilution) for 30 min at room temperature. Cells were then centrifuged at 500 x g for 5 min at 4 °C and washed in FACS buffer. Cells were fixed in 4 % PFA for 15 min at 4 °C, before final wash in FACS buffer and final resuspension in this buffer. Following which, cells were analysed using a BD LSRFortessa X-20 Cell Analyzer (Becton Dickinson). Compensation was done using AbC Total Antibody Compensation Bead Kit (Thermo Fisher Scientific) as per manufacturer’s instructions. A similar but simplified procedure starting with the incubation of cells with the Fc-blocker was performed to evaluate the BOS effect over cell lines and primary cells.

### LDH cell viability assay

As described before [45], post intracellular infection assays, culture supernatants were collected from each well to measure lactate dehydrogenase (LDH) release by using an LDH cytotoxicity assay (Clontech) according to the manufacturer’s instructions. Background LDH activity was determined using mock (PBS)-treated RAW264.7 cells. Maximal LDH activity was determined by lysing cells with 1% Triton X. The percentage of cytotoxicity was calculated as follows: % cytotoxicity = [(sample absorbance − background absorbance)/(maximal absorbance − background absorbance)] × 100.

### Mammalian cell reactive oxygen species quantification

Mammalian cells were seeded at a density of 1×10^6^ in a 6-well tissue culture plate (Black Nunc; Thermo Fisher Scientific), respectively and allowed to attach overnight at 37°C in a 5% CO_2_ humidified atmosphere. BOS treatment was done overnight, and infection with VRE was performed for 3h prior to measuring ROS using the DCFDA / H2DCFDA kit (Abcam) as per the manufacturer’s instructions. TBHP (100 μM) was used as positive control. Plates were incubated with no shaking at 37°C. DHR123 (Sigma) was also used to quantify ROS. Briefly, DHR123 was added to each well to a final concentration of (50 μM) after overnight incubation with BOS (0.52 μg/ml) and/or NAC (5 mM). In the end, cells were detached from the 6-well plate using a cell scrapper and the fluorescence was measured using a BD LSRFortessa X-20 Cell Analyzer (Becton Dickinson) to determine cellular ROS levels.

### RNA isolation, sequencing, and data analysis

RNAs were extracted with RNeasy MinElute Kit (Qiagen), converted into cDNA and sequenced using an Illumina Hiseq®2500 v.2 (Illumina, Singapore), 150 bp paired-end. RNA-seq data were aligned to the mouse genome (version mm10) and raw counts of each gene of each sample were calculated with bowtie2 2.2.3 [46] and RSEM 1.2.1555 [47]. Differential expression analysis was performed using the program edgeR at P < 0.05 with a two fold-change [48]. The gene expression level across different samples was normalized and quantified using the function of cpm. DEGs were annotated using online functional enrichment analysis tool DAVID (http://david.ncifcrf.gov/) [49]. Gene set enrichment analysis were performed with GSEA [50] with FDR q-value < 0.05. Histograms were generated using Prism 9.2.0 (Graphpad).

### NF-κB reporter assay

This assay was performed as described in [45] using RAW-blue cells (InvivoGen). Post treatment of RAW267.4 cells for 15h with BOS (0.52 μg/ml) or LPS (100 ng/mL) and IFN-γ (50 ng/mL) or IL-4 (10 ng/mL) and IL-13 (10 ng/mL) with or without VRE infection, 20 μl of supernatant was added to 180 μl of Quanti-Blue reagent (Invivogen) and incubated overnight at 37°C. SEAP levels were determined at 640 nm by using a Tecan M200 microplate reader.

### Annexin V apoptosis assay

Annexin V apoptosis assay was performed as per manufacturer’s instructions (BD Bioscience). Cells were analysed post infection and BOS treatment within 1h post Annexin V and PI staining.

### Statistical analysis

Statistical analysis was done using Prism 9.2.0 (Graphpad). We used non-parametric Mann-Whitney Test to compare ranks, and one-way analysis of variance (ANOVA) with appropriate post-tests, as indicated in the figure legend for each figure (unless otherwise stated) to analyze experimental data comprising 3 independent biological replicates, where each data point is typically the average of a minimum 2 technical replicates (unless otherwise noted). In all cases, a p-value of ≤0.05 was considered statistically significant.

## Data and materials availability

All data associated with this study are present in the paper or the Supplementary Materials.

## Acknowledgments

We thank Ms Rachel Tan Jing Wen for support with animal work.

## Author contributions

R.A.G.D.S. and C.J.S. performed experiments. R.A.G.D.S., C.J.S. and G.H. analyzed data. R.A.G.D.S., K.A.K. and J.C. interpreted the results. R.A.G.D.S., K.A.K. and J.C. designed the experiments and devised the project. R.A.G.D.S., K.A.K. and J.C. wrote the manuscript. All authors reviewed and approved the final manuscript.

## Supplementary figures

**Figure S1.**
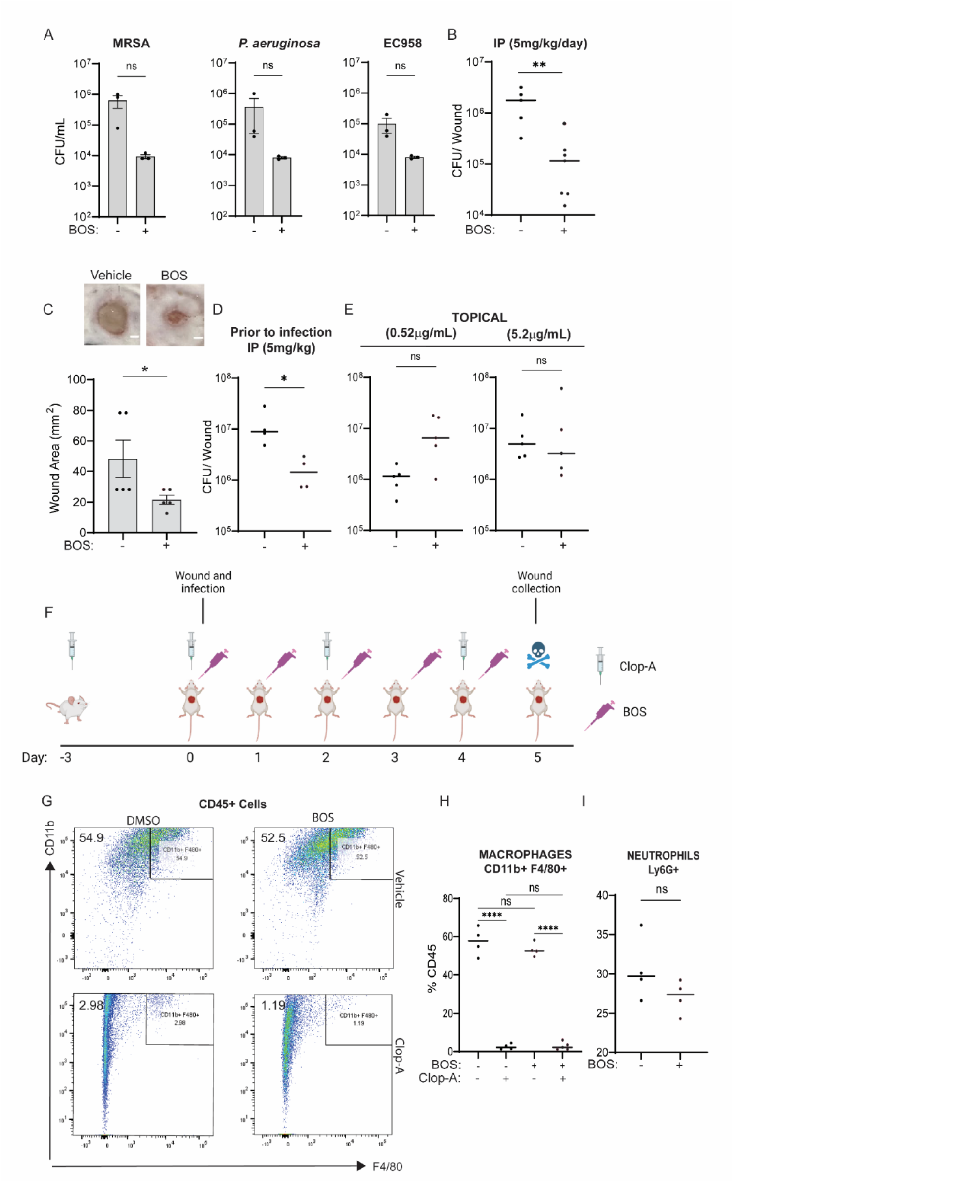
- Macrophages are required for BOS-induced phenotype *in vivo*. **(A)** Comparison of MRSA, *P. aeruginosa* and *E. coli* EC958 CFU in RAW264.7 in the presence or absence of BOS (0.52 μg/mL). **(B)** Comparison of VRE CFU per infected wound of animals treated with five IP injections of vehicle or BOS (30 μL of 5mg/kg). **(C)** Representative images of wounds (top panel) and summary of data from five mice (low panel) at the end of the multiple-treatment experiment. Scale bars, 2 mm. Wound area measured at 4 dpi after five treatments. **(D)** Comparison of VRE CFU per infected wound of animals treated with a single IP injection of vehicle or BOS (5mg/kg) 24h prior to infection. **(E)** Comparison of VRE CFU per infected wound treated topically a single dose of vehicle or BOS (5.20 μg/mL). **(F**-**I)** A schematic diagram of experimental design (F). Mice were injected IP with clop-A (200 μL, 6 mg/mL) 3 days prior to wounding and infection, and additional doses of clop-A on the day of wounding and infection and every 2 days afterwards. In addition, clop-A (10 μL, 6 mg/mL) was applied to the wounds every 2 days. Following VRE infection, BOS (10μL, 0.52 μg/ml) was applied to the wounds daily for 5 days. Five days after wounding and infection, mice were sacrificed, and wounds were recovered for assaying macrophage depletion and VRE CFU. Representative flow cytometry of macrophages (CD45^+^ CD11b^+^ F4/80^+^) from infected wounds (G). The number indicate percentages of cells within the gated areas. Comparison of the percentages of macrophages recovered from infected wounds with or without clop-A and/or BOS treatments (H). Comparison of the percentages of neutrophils recovered from infected wounds that were vehicle or BOS treated with five topical doses (I). Each symbol represents one mouse with the median indicated by the horizontal line (B, D-E, and H-I). Data were from at least two independent experiments with two to three mice per experiment. Statistical analysis was performed using unpaired t test with Welch’s corrections (A), using the nonparametric Mann-Whitney test to compare ranks or using Kruskal-Wallis test with uncorrected Dunn’s posttest (B-E, H-I)). NS, P > 0.05; *P ≤ 0.05, **P ≤ 0.01, ***P ≤ 0.001 and ****P ≤ 0.0001.

**Figure S2.**
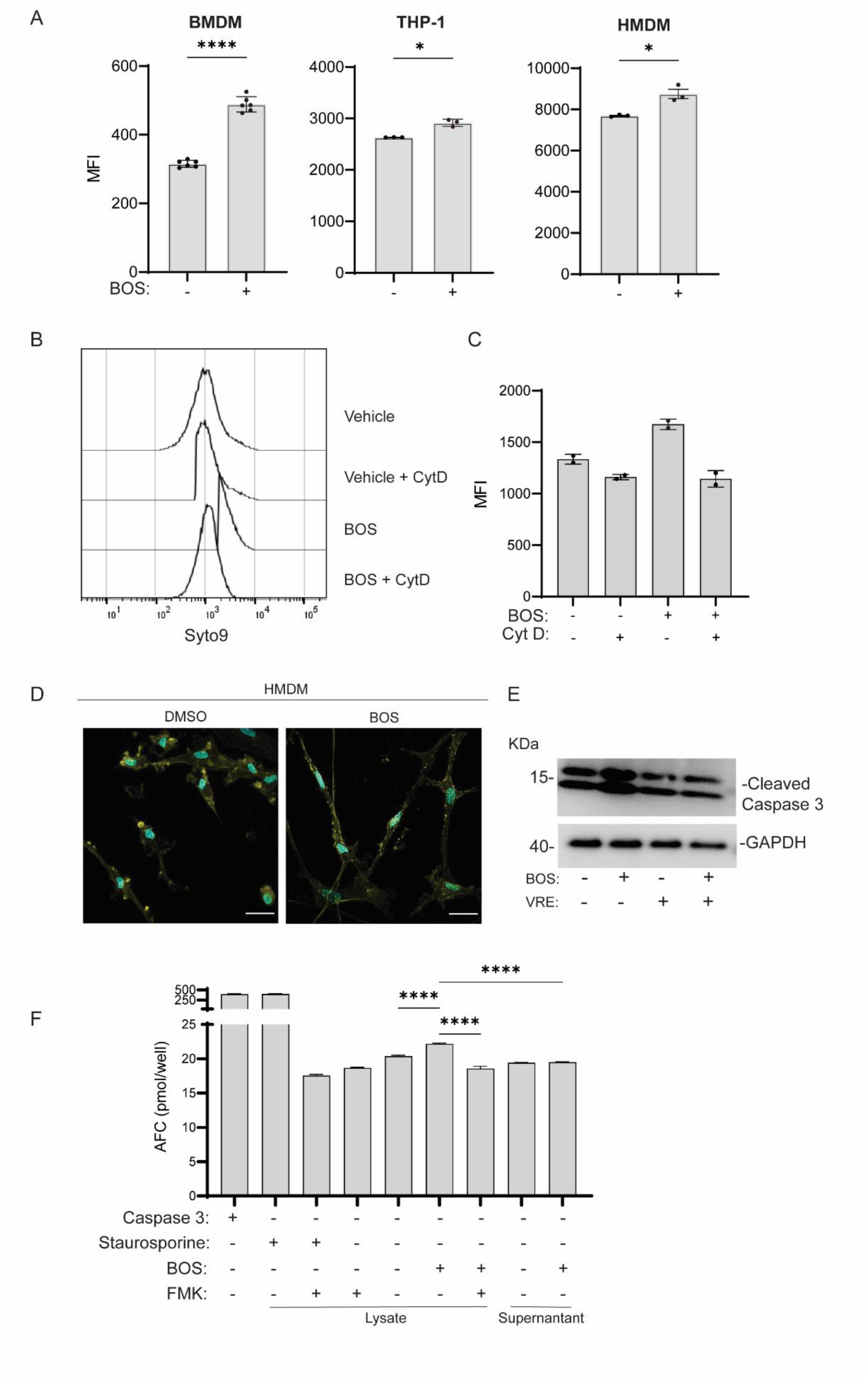
- BOS stimulates macrophage phagocytosis. **(A)** Phagocytosis of VRE by BMDM, THP-1 and HMDM in the presence or absence of BOS. Data (mean ± SEM) are summary of at least three independent experiments. **(B-C)** Comparison of uptake of SYTO9-labelled VRE by RAW264.7 macrophages in the presence or absence of BOS or CytD. RAW264.7 macrophages with and without BOS pre-treatment in the presence or absence of CytD were infected for 1h with SYTO9-labelled VRE, followed by quenching with trypan blue and measurement of fluorescence intensity by flow cytometry. Shown are the representative staining profiles (B) and MFI from two independent experiments (C). **(D)** Representative CLSM images of DMSO or BOS treated HMDM samples that were stained with phalloidin for actin visualization and Hoechst 33342 for nucleus visualization. Images are maximum intensity projections of the optical sections (0.64 μm z-volume) and are representative of 3 independent experiments. Scale bar: 20 µm. **(E)** Western blotting analysis of cleaved caspase 3 in RAW264.7 cells in response to BOS treatment and VRE infection. Whole cell lysate was Western blotted with anti-cleaved caspase 3 and anti-GAPDH. **(F)** Caspase 3 activity assay of cell lysates or supernatants of RAW234.7 cells that were non-treated or treated with BOS. Staurosporine (100 μM) and FMK (50 μM) were also included as positive and negative controls for intracellular Caspase 3 activation, respectively. Data (mean ± SEM) are a summary of at least three independent experiments. Statistical analysis was performed using unpaired t test with Welch’s corrections (A, C) or ordinary one-way ANOVA, followed by Tukey’s multiple comparison test (F). *P ≤ 0.05, and ****P ≤ 0.0001.

**Figure S3.**
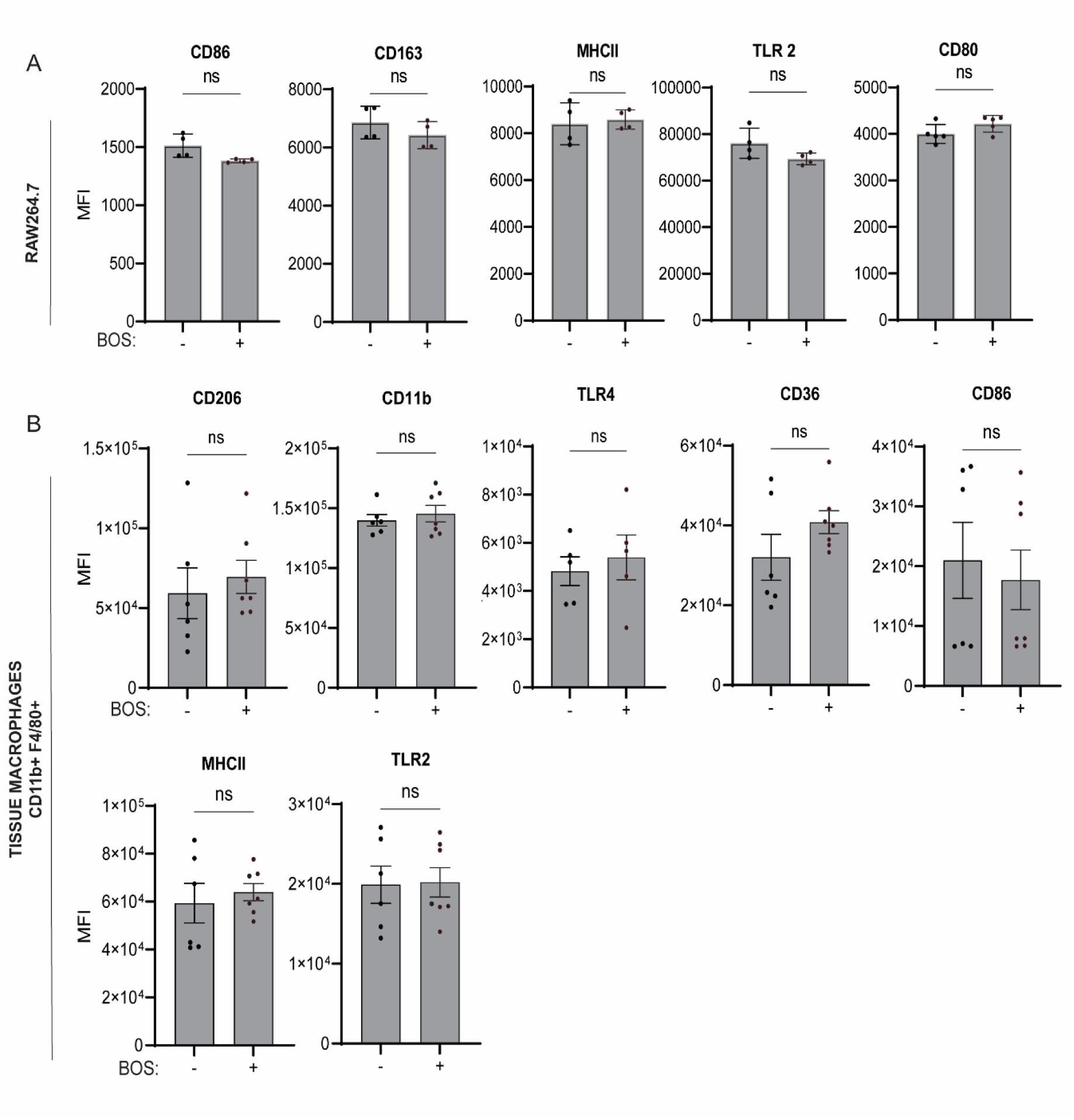
- Effect of BOS treatment on the expression of surface markers related to bacterial uptake. **(A-B)** Comparison of MFI of bacterial recognition, uptake and presentation surface markers gating on CD45^+^ RAW264.7 macrophages non-treated or treated with BOS (A) and CD45^+^ CD11b^+^ F4/80^+^ macrophages from wounds of animals treated with an IP injection of vehicle (-) or BOS (+) (B). Data (mean ± SEM) are summary of at least two independent experiments (A-B) with two to four mice per experiment. Statistical analysis was performed using unpaired t test with Welch’s corrections. NS, P > 0.05; *P ≤ 0.05, **P ≤ 0.01, ***P ≤ 0.001, and ****P ≤ 0.0001.

**Figure S4.**
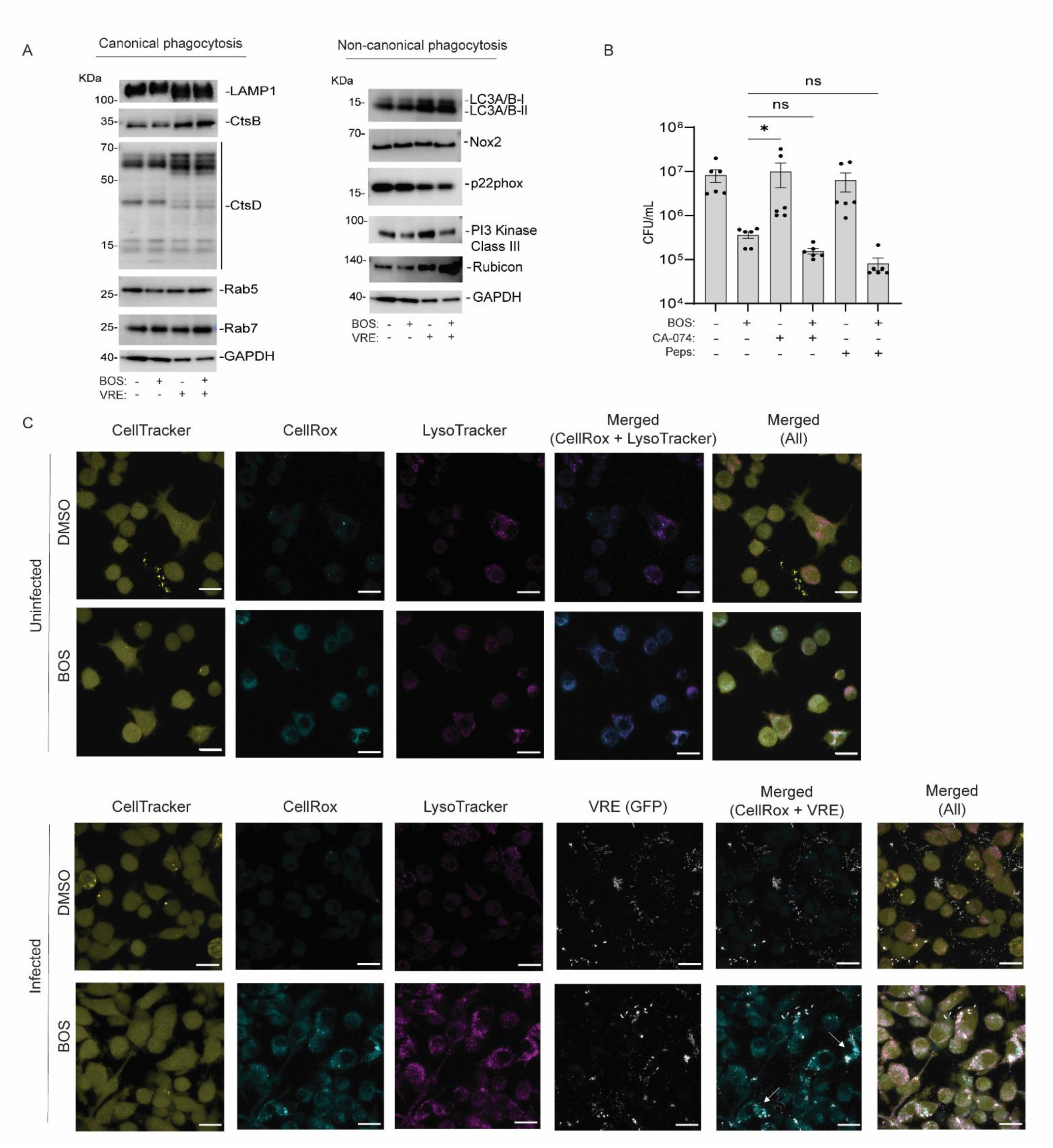
- BOS-treated macrophages produce more ROS. **(A)** Western blot of whole-cell lysates for proteins of the canonical and non-canonical phagocytosis pathway. RAW264.7 cells with (+) and without (−) VRE infection were treated with BOS (+) or left untreated (−). Cell lysates were Western blotted with various antibodies and anti-GAPDH. Shown are representative data from at least two independent experiments. **(B)** Effect of cathepsin inhibitors on BOS-stimulated bacterial killing by macrophages. RAW264.7 cells were infected with VRE in the presence of BOS (0.52 μg/mL), CtsB inhibitor CA-074 (5 nM), and CtsD inhibitor Pepstatin A (Peps, 10 μg/mL) alone or in combination. Intracellular bacterial CFU was quantified after 18h. Data (mean ± SEM) are a summary of at least three independent experiments. Statistical analysis was performed using ordinary one-way ANOVA, followed by Tukey’s multiple comparison test. NS, P > 0.05; *P ≤ 0.05, **P ≤ 0.01, and ****P ≤ 0.0001. **(C)** Visualization of ROS in RAW264.7 cells following BOS and VRE infection by microscopy. Representative CLSM images of DMSO or BOS-treated RAW264.7 samples that were stained with CellTracker (yellow) for cell shape visualization, CellRox (cyan) for ROS visualization and LysoTracker (magenta) for lysosomes visualization. Bottom panels were also infected with pDasher GFP-expressing VRE cells (gray). White arrows point to areas with intracellular VRE cells and high levels of ROS. Images are maximum intensity projections of the optical sections (0.64 μm z-volume) and are representative of at least 2 independent experiments. Scale bar: 20 µm.

**Figure S5.**
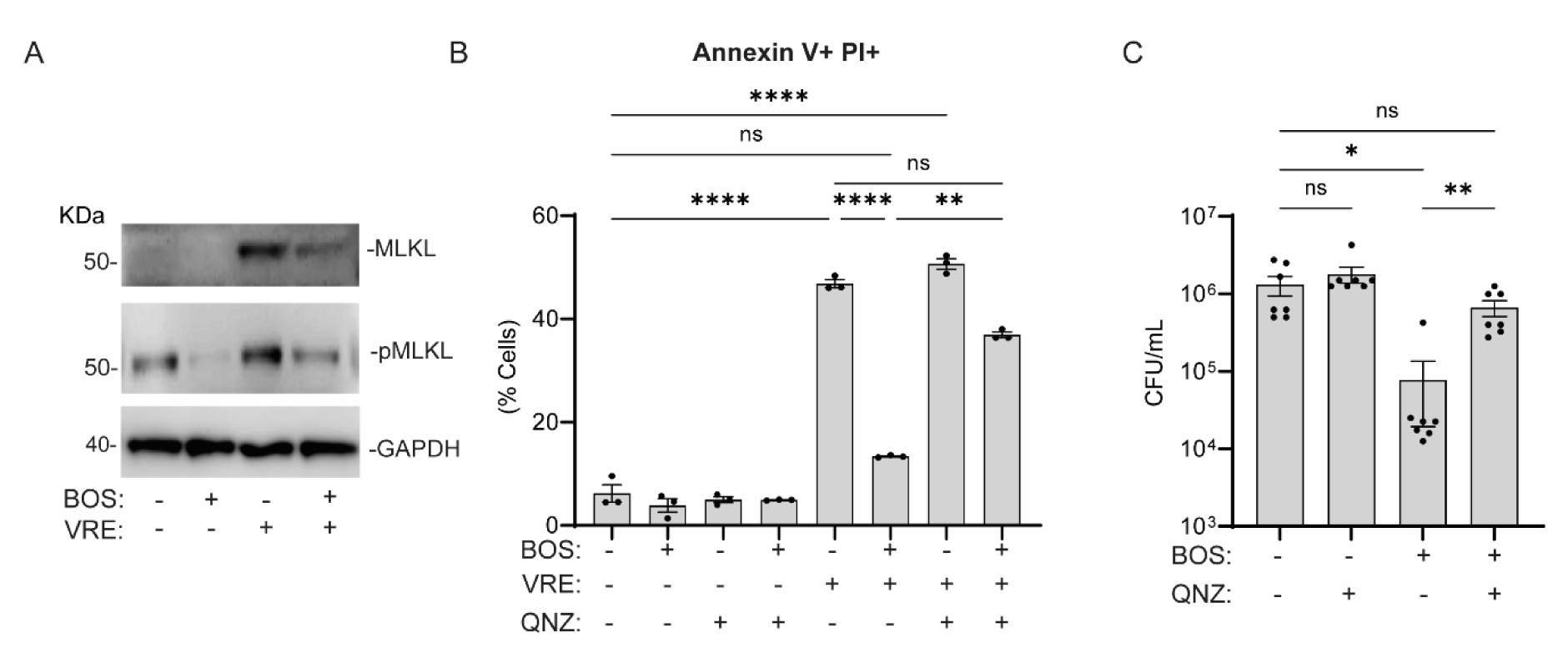
- BOS promotes survival of infected macrophages. **(A)** Western blotting analysis of MLKL and pMLKL. RAW264.7 cells with (+) and without (−) VRE infection were treated with BOS (+) or left untreated (−). Whole-cell lysates were Western blotted with anti-MLKL, anti-pMLKL and anti-GAPDH antibodies. **(B)** Comparison of percentage of Annexin V^+^ and PI^+^ cells at the end of infection. RAW264.7 cells were either infected or not infected and were treated with BOS alone or in combination with the NF-κB inhibitor QNZ (10 nM). Annexin V and PI reactivity was assayed by flow cytometry. Data (mean ± SEM) are a summary of at least three independent experiments. **(C)** RAW264.7 cells were infected with VRE in the presence of BOS (0.52 μg/mL), and QNZ (1 nM), alone or in combination. Intracellular bacterial CFU was quantified after 18 h. Statistical analysis was performed using ordinary one-way ANOVA, followed by Tukey’s multiple comparison test (B), or Brown-Forsythe and Welsh ANOVA test (C). NS, P > 0.05; *P ≤ 0.05, **P ≤ 0.01, and ****P ≤ 0.0001.

## Supplementary Tables

**Table S1.** - Bacterial strains used in this study.

| Strain | Reference |
| --- | --- |
| *E. faecalis* V583 (VRE) | [51] |
| *E. faecalis* V583 + pDasher GFP | This study |
| *S. aureus* USA300 (MRSA) | [52] |
| *E. coli* EC958 | [53] |
| *P. aeruginosa* PAO1 | [54] |

**Table S2.** - MIC of BOS and antibiotics alone or in combination with 0.52 µg/mL BOS.

| Compound or Antibiotic | MIC (µg/mL) | MIC in combination with BOS (0.52 µg/mL) | Strain |
| --- | --- | --- | --- |
| BOS | >13 | - | VRE |
| Vancomycin | 18 | 18 | VRE |
| Penicillin G | 2 | 2 | VRE |
| Gentamycin | 1000 | 500 | VRE |
| BOS | >13 | - | MRSA |
| BOS | >13 | - | *P. aeruginosa* PAO1 |
| BOS | >13 | - | *E. coli* EC958 |

**Table S3.**
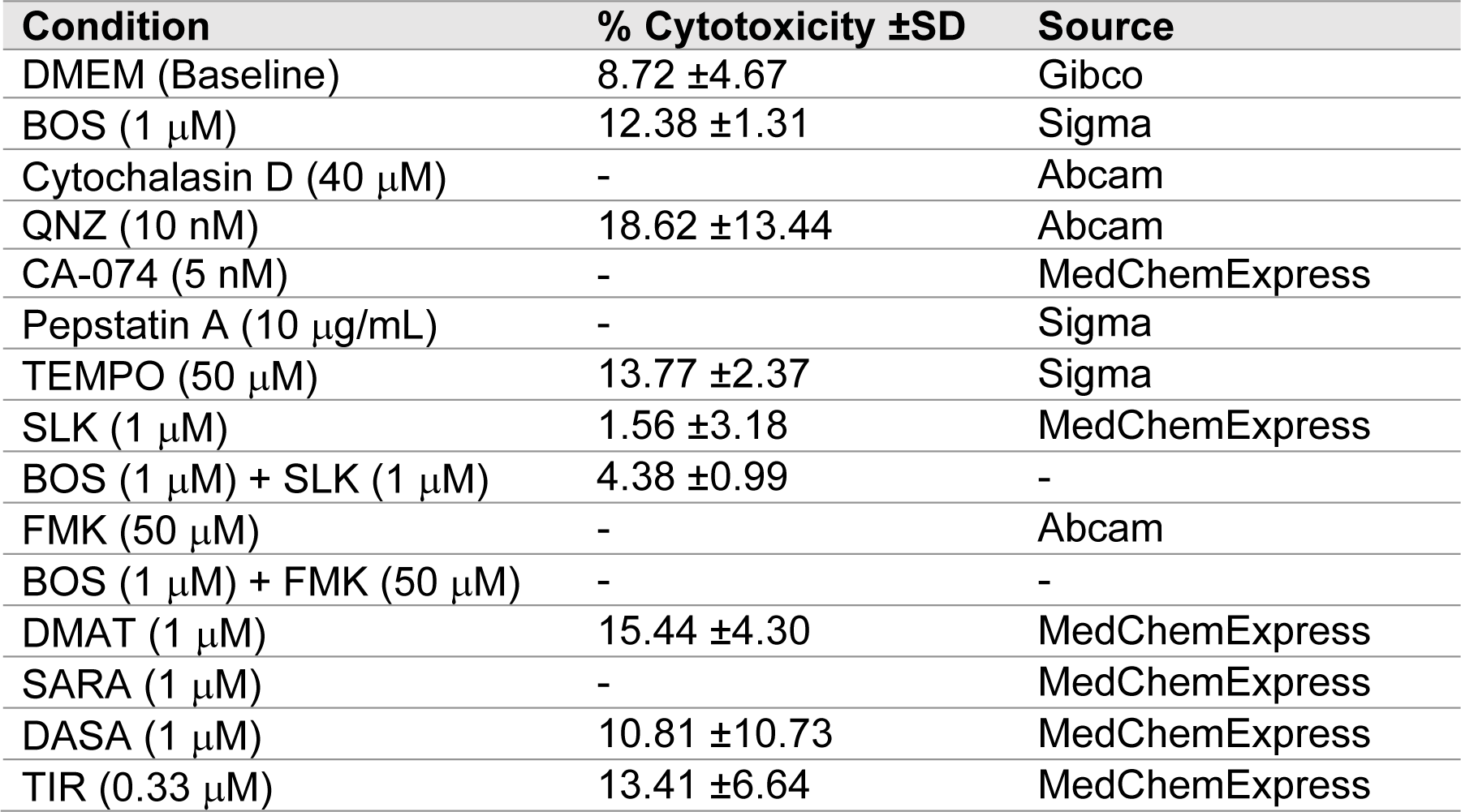
- Cytotoxicity as measured by LDH assay of compounds used in this study.

| Condition | % Cytotoxicity ±SD | Source |
| --- | --- | --- |
| DMEM (Baseline) | 8.72 ±4.67 | Gibco |
| BOS (1 μM) | 12.38 ±1.31 | Sigma |
| Cytochalasin D (40 μM) | - | Abcam |
| QNZ (10 nM) | 18.62 ±13.44 | Abcam |
| CA-074 (5 nM) | - | MedChemExpress |
| Pepstatin A (10 μg/mL) | - | Sigma |
| TEMPO (50 μM) | 13.77 ±2.37 | Sigma |
| SLK (1 μM) | 1.56 ±3.18 | MedChemExpress |
| BOS (1 μM) + SLK (1 μM) | 4.38 ±0.99 | - |
| FMK (50 μM) | - | Abcam |
| BOS (1 μM) + FMK (50 μM) | - | - |
| DMAT (1 μM) | 15.44 ±4.30 | MedChemExpress |
| SARA (1 μM) | - | MedChemExpress |
| DASA (1 μM) | 10.81 ±10.73 | MedChemExpress |
| TIR (0.33 μM) | 13.41 ±6.64 | MedChemExpress |

**Table S4.** - Comparison of transcript levels of cell surface markers associated with bacterial recognition, uptake, and presentation in RAW264.7 cells with and without BOS treatment.

| Gene | logFC | pValue |
| --- | --- | --- |
| *CD36* | -2.5004664 | 1.31E-12 |
| *CD80* | 1.06914694 | 3.32E-06 |
| *CLEC7A*<br>(*DECTIN-1*) | -0.1803542 | 2.79E-01 |
| *CD14* | -0.2248473 | 1.39E-01 |
| *ITGAM (CD11B)* | 0.74697971 | 8.90E-07 |
| *TLR4* | 0.48315495 | 2.53E-04 |
| *TLR2* | -0.0713851 | 5.92E-01 |
| *TLR8* | 1.11318224 | 8.89E-12 |

